# SH3KBP1 scaffolds endoplasmic reticulum and controls skeletal myofibers architecture and integrity

**DOI:** 10.1101/2020.05.04.076208

**Authors:** Alexandre Guiraud, Emilie Christin, Nathalie Couturier, Carole Kretz-Remy, Alexandre Janin, Alireza Ghasemizadeh, Anne-Cécile Durieux, David Arnould, Norma Beatriz Romero, Mai Thao Bui, Vladimir L. Buchman, Laura Julien, Marc Bitoun, Vincent Gache

## Abstract

The building block of skeletal muscle is the multinucleated muscle fiber, formed by the fusion of hundreds of mononucleated precursor cells, myoblasts. In the normal course of muscle fiber development or regeneration, myonuclei are actively positioned throughout muscular development and adopt special localization in mature fibers: regular spacing along muscle fibers periphery, raising the notion of MyoNuclear Domains (MNDs). There is now growing support for a direct connection between myonuclear positioning and normal function of muscles, but how myonuclei affects muscle function remains poorly characterized.

To identify new factors regulating forces applied on myonuclei in muscles fibers, we performed a siRNA screen and identified SH3KBP1 as a new factor controlling myonuclear positioning in early phases of myofibers formation. Depletion of SH3KBP1 induces a reset of MNDs establishment in mature fibers reflected by a dramatic reduction in pairwise distance between myonuclei. We show that SH3KBP1 scaffolds Endoplasmic Reticulum (ER) in myotubes that in turn controls myonuclei velocity and localization and thus myonuclear domains settings. Additionally, we show that in later phases of muscle maturation, SH3KBP1 contributes to the formation and maintenance of Sarcoplasmic Reticulum (SR) and Transverse-tubules (T-tubules). We also demonstrate that in muscle fibers, GTPase dynamin-2 (DNM2) binds to SH3 domains of SH3KBP1. Interestingly, we observed that *Sh3kbp1* mRNA is up regulated in a mouse model harboring the most frequent mutation for Autosomal Dominant CentroNuclear Myopathy (AD-CNM): *Dnm2*^+/R465W^. SH3KBP1 thus appears as a compensation mechanism in this CNM model since its depletion contributes to an increase of CNM-like phenotypes (reduction of muscle fibers Cross-section Areas (CSA) and increase in slow fibers content).

Altogether our results identify SH3KBP1 as a new regulator of myonuclear domains establishment in the early phase of muscle fibers formation through ER scaffolding and later in myofibers integrity through T-tubules scaffolding/maintenance.

**Summary:** Myonuclei are actively positioned throughout muscular development. Guiraud, Christin, Couturier *et al* show that SH3KBP1 scaffolds the ER through Calnexin interaction and controls myonuclei motion during early steps of muscle fibers formation. Besides SH3KBP1 participates in cell fusion and T-tubules formation/maintenance in mature skeletal muscle fibers and contributes to slow-down CNM-like phenotypes.

## Introduction

The building block of skeletal muscle is the multinucleated muscle fiber, formed by the fusion of hundreds of specialized cells, myoblasts, and in which positioning of nuclei (“myonuclei”) is finely regulated. In the normal course of muscle development or regeneration, myonuclei actively localize themselves and adopt specific position in mature myofibers in which they are regularly spaced at the periphery of myofibers (Bruusgaard *et al*, 2003). This precise myonuclei spatial organization gives rise to the notion of MyoNuclear Domains (MNDs) in which each myonucleus controls genes expression in its surrounding cytoplasm and guaranties muscle function (Gundersen, 2016). MNDs settings mainly depend on muscle fibers ability to maintain a defined distance between myonuclei in a cytoplasmic-adapted context related to myofibers type, size and age (Bruusgaard *et al*, 2006; Qaisar & Larsson, 2014; Liu *et al*, 2009). During the first steps of muscle fibers formation, myonuclei have been shown to move and position using mainly microtubule network reorganization (Tassin *et al*, 1985; Gimpel *et al*, 2017) and an interplay between microtubule associated proteins (MAPs) such as MAP7 (Metzger *et al*, 2012) and molecular motors, including dynein and kinesins (Gache *et al*, 2017). This dynamic of myonuclei contributes to precocious alignment of myonuclei in immature fibers. Although comprehension of mechanisms involved in myonuclear positioning during myofibers formation has recently progressed, how mispositioning of myonuclei affects muscle function still remains an open question.

Abnormal myonuclei internalization in muscle fibers that are not directly linked to excessive regenerative process is the hallmark of a group of humans myopathies called Centronuclear Myopathies (CNMs)(Romero & Bitoun, 2011; Jungbluth *et al*, 2007). The majority of defective proteins implicated in CNMs are involved in various aspects of membrane trafficking and remodeling and are relevant to essential cellular processes including endocytosis, intracellular vesicle trafficking and autophagy (Romero & Laporte, 2013). To date, the main proteins implicated in CNMs are phosphatidylinositol phosphatase Myotubularin coded by *MTM1* gene (Laporte *et al*, 1996); GTPase dynamin-2 involved in endocytosis and cell motility, and coded by *DNM2* gene (Bitoun *et al*, 2005); amphiphysin-2 nucleocytoplasmic adaptor protein, coded by *BIN1* gene (Nicot *et al*, 2007) and the principal sarcoplasmic reticulum calcium release channel, ryanodine receptor 1, coded by *RYR1* gene (Wilmshurst *et al*, 2010; Bevilacqua *et al*, 2011).

Myonuclei spatial organization can contribute to muscle fibers functionality as growing evidences support a direct connection between myonuclear positioning and normal function of muscles (Metzger *et al*, 2012; Falcone *et al*, 2014; Janin & Gache, 2018; Robson *et al*, 2016). Additionally, T-tubule organization and efficiency are highly impacted in CNM (Chin *et al*, 2015; Al-Qusairi *et al*, 2009). Finally, besides abnormal nuclear positioning and T-tubules abnormalities, altered autophagy is frequently described in CNM (Jungbluth & Gautel, 2014).

SH3KBP1, also known as Ruk/CIN85 (Cbl-interacting protein of 85 kDa) is a ubiquitously expressed adaptor protein, involved in multiple cellular processes including signal transduction, vesicle-mediated transport and cytoskeleton remodeling (Havrylov *et al*, 2010; Buchman *et al*, 2002). SH3KBP1 protein is composed of three SH3 domains at the N-terminus followed by a proline-rich (PR) domain, a serine-rich (SR) domain, and a C-terminus coiled-coil (CC) domain. Functions of SH3KBP1 as an adaptor protein are mainly linked to endocytosis trafficking and degradative pathway through the recruitment of Cbl (E3 ubiquitin ligase), endophilin and dynamin (Schroeder *et al*, 2010; Sun *et al*, 2015; Zhang *et al*, 2009). SH3KBP1 protein is also associated with several compartments involved in membrane trafficking such as the Golgi complex and is mainly concentrated in COPI-positive subdomains (Havrylov *et al*, 2008). The role of SH3KBP1 during the formation of muscle fibers has never been investigated.

In the present study, we show that the adaptor protein SH3KBP1, through its N-terminus part, is able to scaffold endoplasmic reticulum (ER) probably *via* an interaction with Calnexin. SH3KBP1 also governs myonuclei motion and spatial organization as well as myoblast fusion. Additionally, we observed that SH3KBP1 interacts with Dynamin 2 (DNM2) and organizes T-tubule formation in skeletal muscle. Moreover, its down-regulation contributes to an increase of CNM-like phenotypes in a mouse model expressing the most frequent mutation causing autosomal dominant centronuclear myopathy (AD-CNM).

## Results

### SH3KBP1 is required for precocious myonuclear positioning steps and controls fusion parameters during muscle fiber formation

To identify new factors that contribute to myonuclear spreading and alignment in muscle fibers, we performed a large siRNA screen on candidates expected to affect myonuclei repartition in nascent myotubes. Briefly, primary myoblasts were isolated from young pups and induced to differentiate *in vitro* for 3 days before analysis (Falcone *et al*, 2014) (Fig.1A). This cell culture technique leads to an accumulation of heterogeneous myotubes regarding myonuclei number and myotubes length (Blondelle *et al*, 2015). To precisely characterize the early steps of myonuclei positioning in myotubes, we developed an image-J^®^ plugin that automatically extracts different myotubes parameters (such as myotubes lengths/areas and myonucleus localization). Data were then classified and analyzed according to myonuclei content using a homemade program in R-Studio^®^ (Fig.1D-F). This analysis allowed us to track myonuclei accretion during elongation of myotubes in a window from 3 to 11 myonuclei per myotubes. Isolated murine myoblasts were then treated with short interfering RNA (siRNA) targeting either a scrambled sequence or candidate genes. Among candidates from the siRNA screen, efficient *sh3kbp1* depletion appears to strongly modify myonuclear positioning (Fig1. B-C, Fig.S1A). In control condition, we observed a nearly linear relation between addition of myonuclei into myotubes and expansion of myotubes length, with an average of nearly 10 % increase of length after each myonucleus accretion (Fig. 1D). In *sh3kbp1* depleted myotubes, length repartition is homogenously extended to an increase of 46.8% ±3.3 in myotubes containing up to 11 myonuclei (Fig. 1D). Quantification of the mean distance between all myonuclei inside myotubes indicate a burst of spacing (+103.5% ±11.6, data not shown) concomitantly with an escape from the center of myotubes reflected by (i) the mean distance between myotube centroid and each myonuclei (DMcM) (+111.5% ±12.5) (Fig. 1E) and (ii) the Myonuclei Spreading Graphic (MSG) representation (Fig. 1F). MSG shows statistical probability to find one myonucleus along the all length of myotubes and allow the extraction of “statistical clustering zones” (colors code in Fig. 1F) that we estimate as four zones in the case of scramble myotubes *versus* eight zones in *sh3kbp1* depleted myotubes (Fig. 1F). Thus, *sh3kbp1* depleted myotubes failed to homogeneously spread myonuclei along myotubes length that tends to accumulate at myotubes tips (Fig. 1C, arrows). Overall, these data suggest that for a defined myonuclei content in myotubes, SH3KBP1 acts as “anti-elongation” factor and contributes to myonuclei spreading in myotubes.

**Figure 1:**
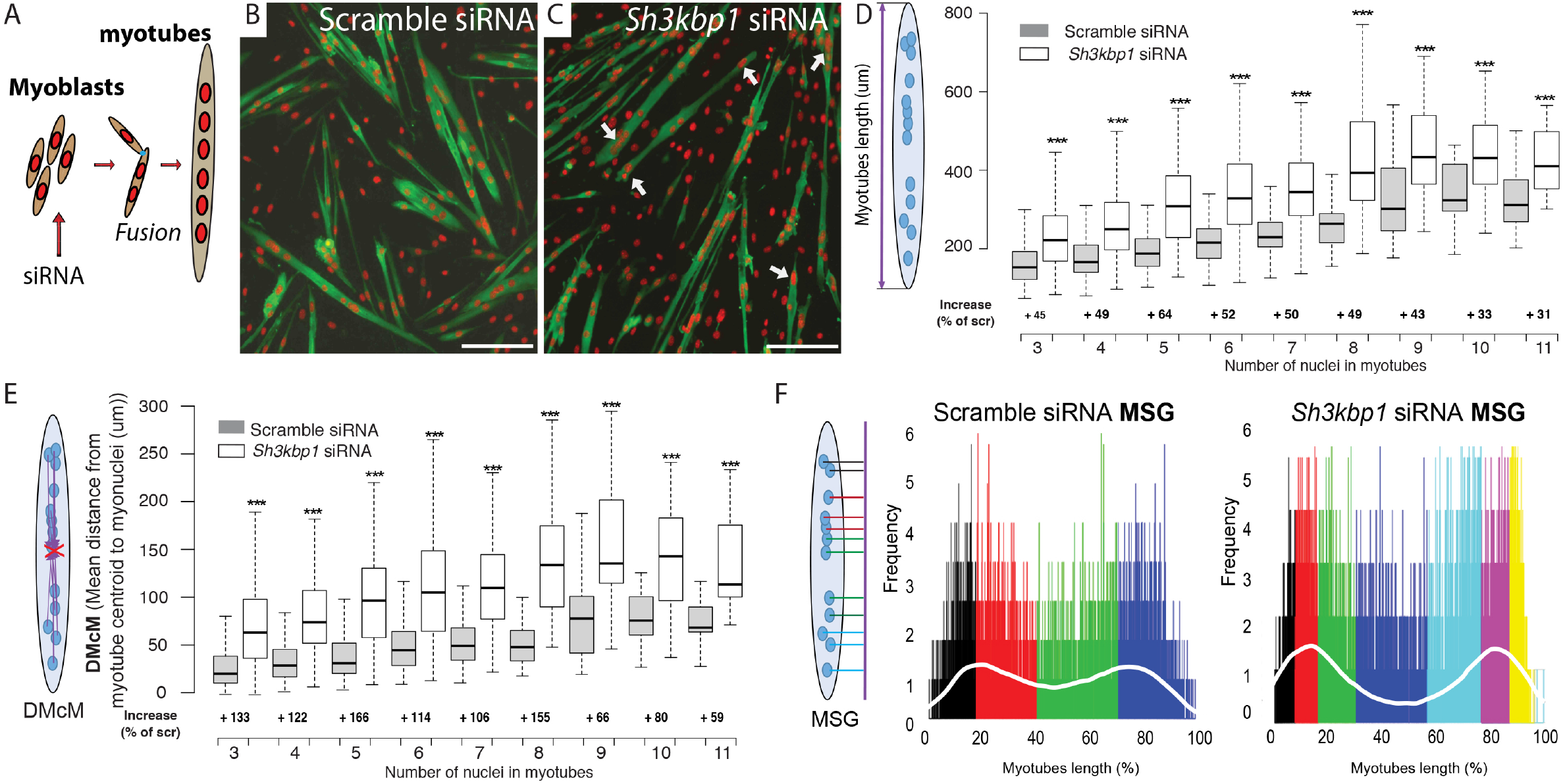
Myonuclear positioning during primary myotubes formation is controlled by *sh3kbp1*. (A) Scheme of sequential steps in order to obtain immature myotubes from primary myoblasts. siRNAs were transfected 24 hours before myoblasts fusion. (B-C) Representative immunofluorescence staining of myosin heavy chain (green) and myonuclei (red) in primary myotubes treated for scramble (B) or a pool of 2 individual siRNAs targeting *Sh3kbp1* mRNA (C) after 3 days of differentiation, white arrows indicate “tips-aggregated myonuclei”. Scale Bar: 150μm. (D-E) Myotubes lengths and Mean distances between each myonuclei and myotube’s centroid (DMcM) ranked by myonuclei content per myotubes were quantified after 3 days of differentiation in cells treated with a scramble or *Sh3kbp1* siRNAs. Data from three independent experiments were combined. Scramble siRNA cells (n=1010) and *Sh3kbp*1 siRNA cells (n=1093), Unpaired t-test, ***p < 0.001. Center lines show the medians; box limits indicate the 25th and 75th percentiles as determined by R software; whiskers extend 1.5 times the interquartile range from the 25th and 75th percentiles, outliers are represented by dots. (F) Myonuclei Spreading Graph (MSG) represents statistical spatial distribution of myonuclei along myotubes in scramble and in *Sh3kbp1* siRNA-treated myotubes; white-line represents the mean value of the statistical frequency.

Myoblasts/myotubes elongation and alignment is a key process that controls myoblast/myotubes fusion (Louis *et al*, 2008). As *sh3kbp1* depletion led to myotubes length increase, we wondered if fusion aspects were modified during differentiation. Indeed, myotubes elongation can contribute to increase myotubes/myoblast contacts that thus will modulate membrane fusion and ultimately modify myonuclei accretion speed (Kim *et al*, 2015). We evaluated fusion capacity of primary myoblasts isolated from young pups after *in vitro* differentiation induction (Fig.S1B-G). After 3 days of differentiation, isolated myoblasts treated with siRNA targeting *sh3kbp1* mRNA are similarly stained for myosin heavy chain antibodies compare to control cells treated with siRNA scramble sequences, indicating normal myoblast commitment into muscle differentiation process (Fig. S1B-D). Fusion index was increased in *Sh3kbp1* depleted myotubes, assessed by the total number of myonuclei in myotubes (Fig. S2E) and the average number of myonuclei per myotube (Fig. S2F). Accordingly, in *sh3kbp1* depleted conditions, distribution of myotubes with respect to myonuclei content revealed much more myotubes with ≥10 nuclei when compared to controls (Fig. S2G). Altogether, this data suggests that *sh3kbp1* acts also as an “anti-fusion” factor as expected for an anti-elongation factor.

### SH3KBP1 governs myonuclei velocity that contributes to myonuclei positioning in mature myofibers

To investigate the role of *sh3kbp1* in late steps of differentiation, primary mouse myotubes were maintained in differentiation media for 5 to 10 days as previously described (Pimentel *et al*, 2017; Falcone *et al*, 2014) Fig. 2A. In 5-day differentiated myofibers, myonuclei clustering and velocity was addressed (Fig. 2B-F). First, we observed twice more myonuclei clustered in myotubes treated with shRNA targeting *sh3kbp1* gene compared to control, confirming a role of *sh3kbp1* on myonuclei spreading (Fig. 2B & E, white asterisk). As myonuclei within myofibers have different behaviors (Gache *et al*, 2017), we next investigated the impact of *sh3kbp1* depletion on myonuclei movements in 5-day differentiated myofibers. Myoblasts were transfected with RFP-lamin-chromobody^®^ to visualize myonuclei concomitantly with shRNA targeting scramble or *Sh3kbp1*, both GFP-tagged. Myotubes containing both constructions (GFP and RFP-lamin-chromobody^®^) were selected for fewer analyses (Fig.2C-F). After 5 days of differentiation, myonuclei were tracked every 20 minutes for a time period of 16 hours (Fig. 2E, Supplementary video 1-2). Myonuclei displacements parameters were analyzed using SkyPad method (Cadot *et al*, 2014). We found that in control condition, myonuclei are in motion during nearly 35 % of the time (a movement is defined as a displacement more than 30 μm) at a median speed of 0.232 ±0,014 μm/min. Depletion of *sh3kbp1* increased from more than 20% the percentage of time myonuclei are in motion of and by more than 30% median speed that reach 0.313 ±0,014 μm/min outside myonuclei clusters (Fig.2C-D). Indeed, because of high myonuclei concentration inside myonuclei clusters, we could not technically access to myonuclei motion and speed (Fig. 2E, Supplementary video 2). To further investigate the implication of *sh3kbp1* in late steps of differentiation, primary mouse myotubes were maintained in differentiation media for 10 days as previously described (Pimentel et al., 2017) (Fig. 2G). In these maturation conditions, myonuclei are compressed between myofiber plasma membrane and contractile apparatus and adopt a flatten architecture all along myofibers length (Roman et al., 2017). This long-term differentiation approach allow to confirm that *sh3kbp1* controls myonuclear positioning during maturation of myofibers as *sh3kbp1* depletion using either siRNA or shRNA caused a significant reduction in the mean distance between adjacent myonuclei (Fig. 2G-H). Additionally, *Sh3kbp1* depletion significantly reduces by more than 30 % myofibers width (Fig. 2I). These results suggest that SH3KBP1 contributes to nuclear spreading and to the reduction of myonuclei movements and motion during muscle fibers maturation.

**Figure 2:**
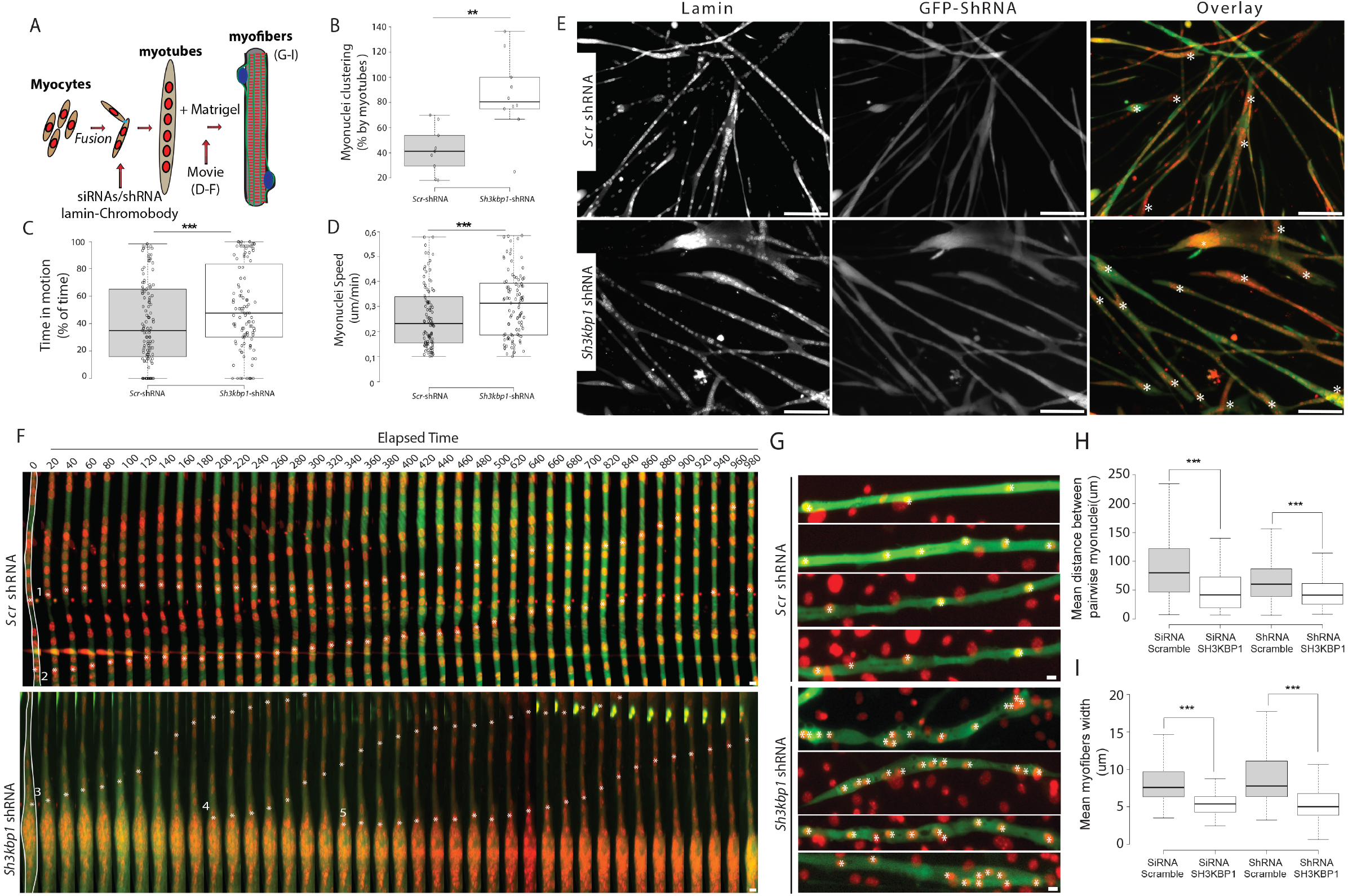
Myonuclei localization and motion are affected in *sh3kbp1* depleted primaries myofibers and lead to myonuclei aggregation. (A) Scheme of sequential steps in order to obtain mature myofibers from primary mice myoblasts. siRNAs or shRNA were transfected during precocious steps of myoblasts fusion. (B-F) Primary myoblasts were treated with scramble or *Sh3kbp1* shRNA tagged with GFP (Green) and co-transfected with RFP-lamin-chromobody^®^ (red) and were induced to differentiate for 5 days into myofibers. (B-D) Quantification of myonuclei clustering by myotubes (B), myonuclei time in motion (C) and myonuclei speed (D) were quantified using SkyPad analysis (Cadot *et al*, 2014). Data from three independent experiments were combined. Unpaired t-test, ***p < 0.001, **p < 0.01. (E) Representative immunofluorescence staining of RFP-lamin-chromobody® (red) and shRNA tagged with GFP (Green) in primary myotubes treated with scramble or *Sh3kbp1* shRNA after 5 days of differentiation into myofibers. Scale Bar: 150μm. (Asterisks represents myonuclei clustering) (F) Frames from a 16h time-lapse movie in two channels (shRNA in green and lamin-chromobody® in red) of primary myotubes. In the first frame (on the left), myofibers are selected in white, which corresponds to the region used to create the adjacent kymograph. Scale Bar: 30μm. (Asterisks 1-5 are examples of individual myonuclei tracking). (G) Four representative images of 10 days differentiated myofibers transfected with either scramble or *Sh3kbp1* shRNA tagged with GFP, shRNA (Green) myonuclei (red). Scale Bar: 10μm. (Asterisks are individual myonuclei) (H-I) Quantification of the mean distance between pairwise myonuclei (H) and mean myofibers width (I) in 10 days differentiated myofibers treated with scramble siRNA or shRNA, a pool of 2 individual siRNAs or an individual shRNAs targeting *Sh3kbp1*. Data from three independent experiments were combined. Center lines show the medians; box limits indicate the 25th and 75th percentiles as determined by R software; whiskers extend 1.5 times the interquartile range from the 25th and 75th percentiles, outliers are represented by dots. Unpaired t-test, ***p < 0.001, **p < 0.01, *p < 0.05.

Next, we focused on the expression of both SH3KBP1 protein and *Sh3kbp1* mRNA in the time course of *in vitro* myotubes formation using C2C12 cells. These analyses showed a slight enhancement of SH3KBP1 production during early stages of myotubes formation (Fig. 3A). These results were confirmed using RT-qPCR techniques where a two-fold increase was observed in mRNA expression at the onset of differentiation step (Fig3B-C). To confirm *Sh3kbp1* functions during muscle cell differentiation, we stably knocked-down *Sh3kbp1* mRNA expression in C2C12 cells using a small hairpin interfering RNA (shRNA) (Fig. 3D). As observed in primary cells, after 3 days of differentiation, myoblasts entered the “muscle differentiation program”, illustrated by the detection of myosin heavy chain-positive (MHC+) cells (Fig. 3E, day3). Although intensity of staining is reduced, we do not observe any alteration of myotubes size repartition in *Sh3kbp1* knocked-down compared to control conditions (Fig. S2A). After 6 days of differentiation, we clearly see a breaking event in fusion capacity correlated with *Sh3kbp1* knockdown (Fig. 3E). In control conditions, myonuclei spread along thin myotubes length while *Sh3kbp1* knockdown gives rise to huge myotubes with clustered myonuclei areas (Fig. 3E, day6). Distribution of myotubes with respect to their myonuclei content reveals also a significant increase in the number of myotubes with high myonuclei content compared to controls (Fig. S2B). In control condition, we observed a limited accumulation of clustered myonuclei along myotubes length, with a majority of clusters containing 4 to 6 myonuclei (Fig. 3F). On the opposite, in *Sh3kbp1* knockdown, we observe a clear increase in the proportion of clusters containing more than 15 myonuclei (Fig. 3F). To address the specificity of myonuclei clustering phenotype in *Sh3kbp1* depleted conditions, we re-expressed full-length SH3KBP1 proteins in *Sh3kbp1* depleted murine myotubes and show that we rescue even better than in control the number of clustered myonuclei by myotubes (Fig. S2C-D). Together, these results show that *Sh3kbp1* is essential during myotube formation, both in primary myoblasts and in C2C12 cell cultures, to control myoblasts fusion and myonuclear positioning during skeletal muscle formation.

**Figure 3:**
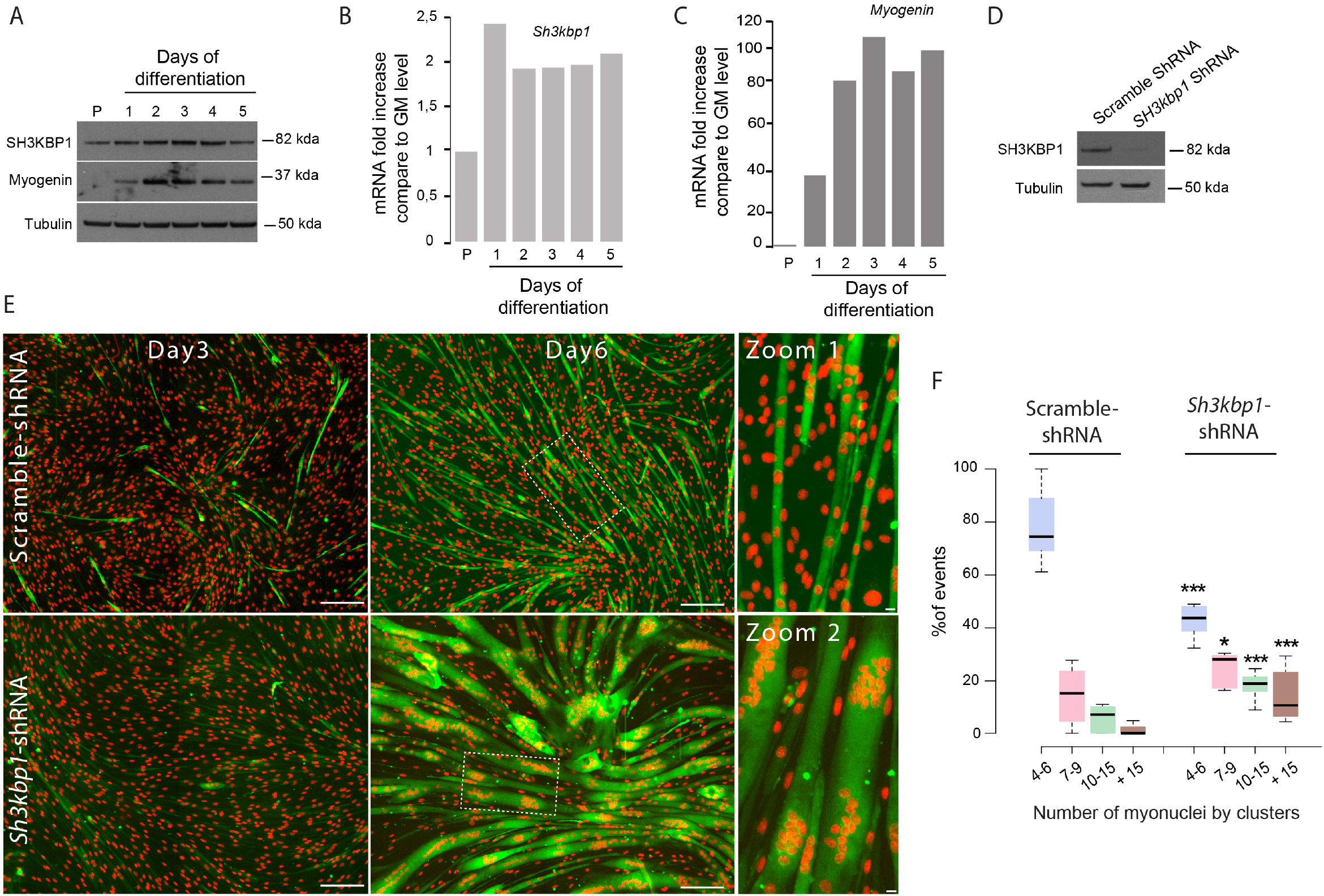
SH3KBP1 is induced during myotubes differentiation and controls myonuclei aggregation in C2C12 cells. (A) Western blot analysis of SH3KBP1 protein expression in total protein extracts from proliferating (P) or differentiating C2C12 cells for up to 5 days. C2C12 differentiation is assessed by Myogenin expression and Tubulin is used as loading control. (B-C) qRT-PCR analysis of *Sh3kbp1* (B) and *Myogenin* (C) gene expression level relative to *CycloB, Gapdh, GusB, Rpl41* and *Tbp* gene in proliferating C2C12 cells (P) or differentiating C2C12 cells for up to 5 days. (D) Representative western blots analysis of SH3KBP1 protein down-regulation in stable cell line constitutively expressing shRNA construct targeting *Sh3kbp1.* Tubulin is used as loading control. (E) Representative immunofluorescence staining of myosin heavy chain (green) and myonuclei (red) in 3 or 6 days cultured C2C12 stable cell lines expressing scramble shRNA or shRNA targeting *Sh3kbp1.* Zooms 1 & 2 are magnifications of images in white dots rectangle in 6 days old myotubes. Scale bars, 150 μm and 15 μm in zoom. (F) Quantification of the number of myonuclei by clusters in C2C12 cells stable cell line expressing scramble shRNA or shRNA targeting *Sh3kbp1* after 5 days of differentiation. Data from three independent experiments were combined. Unpaired t-test, ***p < 0.001, *p < 0.05. Center lines show the medians; box limits indicate the 25th and 75th percentiles as determined by R software; whiskers extend 1.5 times the interquartile range from the 25th and 75th percentiles, outliers are represented by dots.

### SH3KBP1 is an endoplasmic reticulum scaffolding protein that interacts with ER72 and Calnexin

Several studies show that SH3KBP1 is distributed between different membrane trafficking compartments such as the Golgi Complex (Havrylov *et al*, 2008). The three SH3 domains localized in the N-terminal part of SH3KBP1 are responsible for its high capacity to interact with diverse regulatory partners (Havrylov *et al*, 2009) while the C-terminus part, containing a coiled-coil domain allows its targeting to endosomal membranes (Zhang *et al*, 2009). As no role of SH3KBP1 was previously described in skeletal muscle, we analyzed its localization during the time course of muscle formation using both C2C12 myoblast cell line and murine primary myoblasts (Fig. 4 A-B). In growing conditions, SH3KBP1 is localized in the cytoplasm with an apparent higher concentration at the vicinity of nuclei both in primary and in C2C12 myoblasts and some dots inside nucleus are also visible (Fig. 4A-B). During early myotubes formation, SH3KBP1 seems to spread along myotubes length, exhibit a weak perinuclear accumulation and myonuclei still exhibits dots inside myonuclei (Fig. 4A-B, Day3 and Day2). In 5 days differentiation C2C12 myotubes, perinuclear accumulation of SH3KBP1 is stronger and we still observe some accumulation inside some myonucleus as dots or line (Fig. 4A, Day5). In primary myoblasts induced to differentiate into “mature-like” fibers organized with myonuclei at periphery, SH3KBP1 was still detected at myonuclei vicinity but more accumulated at the interface between myonuclei “bottom” and muscle fiber interior and it also exhibited longitudinal/transversal staining (Fig. 4B, Day10). We thus wondered what kind of compartments could be controlled by SH3KBP1. To answer this question, we used stable C2C12 cell-line depleted for *Sh3kbp1* to investigate altered compartment phenotypes (Fig. 4C-D). We observed that, even if myonuclei are clustered in myotubes depleted for *Sh3kbp1*, Golgi marker RCAS1 was still mainly localized around myonuclei, as previously described for Golgi elements (Fig. 4C)(Ralston *et al*, 1999). On the contrary, the ER marker ERP72 that showed perinuclear localization in control myotubes was completely dispersed in *Sh3kbp1*-*depleted* myotubes (Fig. 4D). This result indicates a role for *Sh3kbp1* in the upkeep of ERP72-containing ER specifically at the vicinity of myonuclei, independently of Golgi complex architecture.

**Figure 4:**
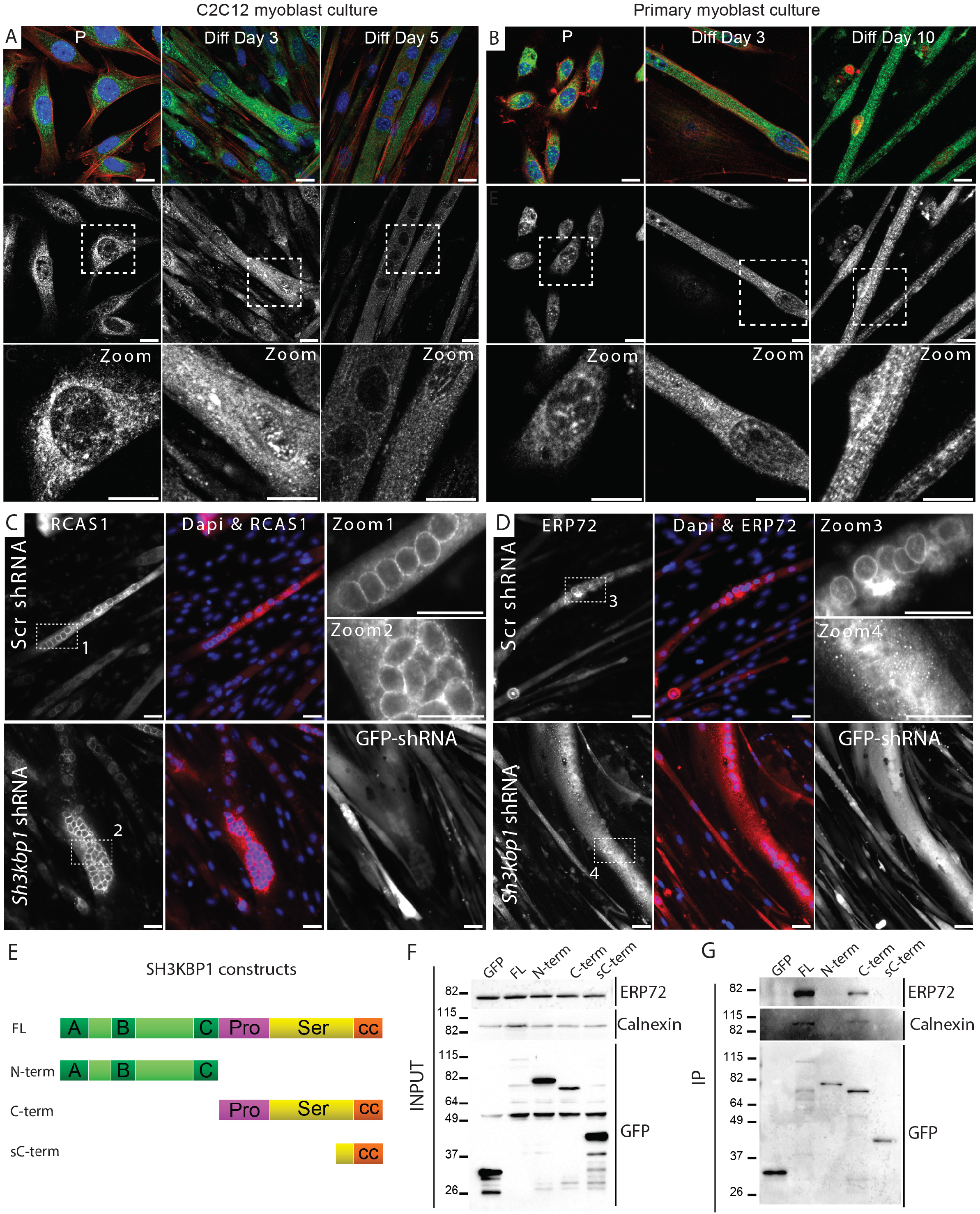
SH3KBP1 is localized near and inside myonuclei and controls ERP72 positives Endoplasmic reticulum through Calnexin interaction during *in vitro* muscle fibers differentiation. (A-B) Representative immunofluorescent staining of SH3KBP1 (green), Actin (Red) and myonuclei (blue) in the course of myotube formation in C2C12 cells line in proliferation (P) or after 2 and 5 days of differentiation (A) or in the course of myofibers formation in primary myoblast cultures in proliferation (P) or after 3 and 10 days of differentiation (B). Scale bars, 10 μm. (C-D) Representative Immunofluorescent staining of Golgi Marker (RCAS1, for receptor binding cancer antigen expressed on SiSo cells, red) (C) and Endoplasmic Reticulum (ERP72, for Endoplasmic reticulum resident protein 72, red) (D) and myonuclei (blue) in 6 days cultured C2C12 stable cell line expressing either scramble shRNA or shRNA targeting *Sh3kbp1* gene (GFP-shRNA). Scale bars, 100 μm. (E) Scheme of SH3KBP1 constructs, labeled with GFP in the N-terminus part, used for immunoprecipitation assay in (F-G). FL: full length. (F) Representative western blot with indicated antibodies (right) in C2C12s expressing GFP-SH3KBP1 constructs (top). (G) Representative western blot with indicated antibodies (right) of GFP-SH3KBP1 constructs immunoprecipitation using C2C12 expressing indicated constructs (top). Blots were repeated more than 3 times.

ERP72 (also called PDIA4) is a disulfide isomerase that acts as a folding chaperone for newly synthesized secretory proteins in the ER compartment (Satoh *et al*, 2005). Endogenous ERP72 co-immunoprecipitated with exogenous full-length GFP-tagged SH3KBP1 (Fig. 4E-G). Using a panel of SH3KBP1 deletion mutants, we determined that ERP72 interacts with the C-terminal part of SH3KBP1, independently of its C-terminus coiled-coil domain (Fig. 4E-G). ERP72 is described as an intraluminal ER protein that interacts with calnexin ER-chaperone (Penga *et al*, 2014). Interestingly, the C-terminus part of Calnexin is cytosolic and thus could be the cytoplasmic linker between SH3KBP1 and ERP72-containing Endoplasmic Reticulum (Wada *et al*, 1991). We next tested if SH3KBP1 physically interacts with Calnexin (Fig. 4E-G) and observed that endogenous Calnexin, as ERP72, co-immunoprecipitated with exogenous full-length GFP-tagged SH3KBP1 constructs. In addition, we also identified the C-terminal part of SH3KBP1, independently of the C-terminus coiled-coil domain as the Calnexin-interacting domain (Fig. 4E-G). Therefore, Proline-and Serine-Rich domains of SH3KBP1 mediate the interaction with ERP72-positive ER through Calnexin binding.

### SH3KBP1 progressively accumulates at the Z-line and interacts with DNM2

We next investigated SH3KBP1 localization *in vivo*, in mature myofibers from adult skeletal muscle. SH3KBP1 accumulated specifically at the vicinity of myonuclei, forming a “cage” around myonuclei in *Tibialis Anterior* (TA) muscle (Fig. 5A-B, asterisks). This staining was confirmed in human muscle biopsies where we observed that myonuclei are positive for SH3KBP1 (Fig. S3G). In TA longitudinal section, SH3KBP1 exhibited a striated pattern at the I-band/Z-line zone, in between the staining of the voltage-dependent calcium channel, DHPRα, which labels T-tubules and forms a doublet band indicating that SH3KBP1 does not colocalize with DHPRα but follow, in a close proximity, T-tubules structures (Fig. 5C-E). To determine domains of SH3KBP1 responsible for this particular localization, we next tested *in vitro* expression of full-length SH3KBP1 or associated fragments into primary myoblasts induced to differentiate into “mature-like” fibers with peripheral myonuclei and proper sarcomere organization, reflected by striated actin staining (Fig. 5F-G). In this myofibers, full-length SH3KBP1 was present as small aggregation patches close to myonuclei, combined with striated pattern and accumulated at the Z-line with no overlap with actin staining (Fig. 5F-G, SH3KBP1-FL). N-terminus fragment of SH3KBP1 containing SH3 domains was only present as striated patterns with no particular accumulation at myonuclei vicinity, while the C-terminus SH3KBP1 fragments strongly accumulated at the periphery of muscle fibers, at the vicinity of myonuclei and at the Z-line with striated patterns (Fig. 5.F). Numerous proteins accumulate at the I-band/Z-line zone and participate in myofibers structuration (Burgoyne *et al*, 2015). Among them, Dynamin 2 (DNM2), a large GTPase implicated in cytoskeleton regulation and endocytosis, was previously described in HeLa cells as a SH3KBP1 interacting protein, through its proline-rich domain (Schroeder *et al*, 2010). To this end, we expressed, in C2C12 cells, full-length GFP-tagged-DNM2 with fragments of FLAG-tagged SH3KBP1 and immunoprecipitated SH3KBP1 constructs. This experiment confirmed that DNM2 interacts with SH3KBP1 through its N-terminal part (Fig. 5H-I). Therefore, in mature myotubes, SH3KBP1 accumulate at the Z-line where it forms a protein complex with DNM2.

**Figure 5:**
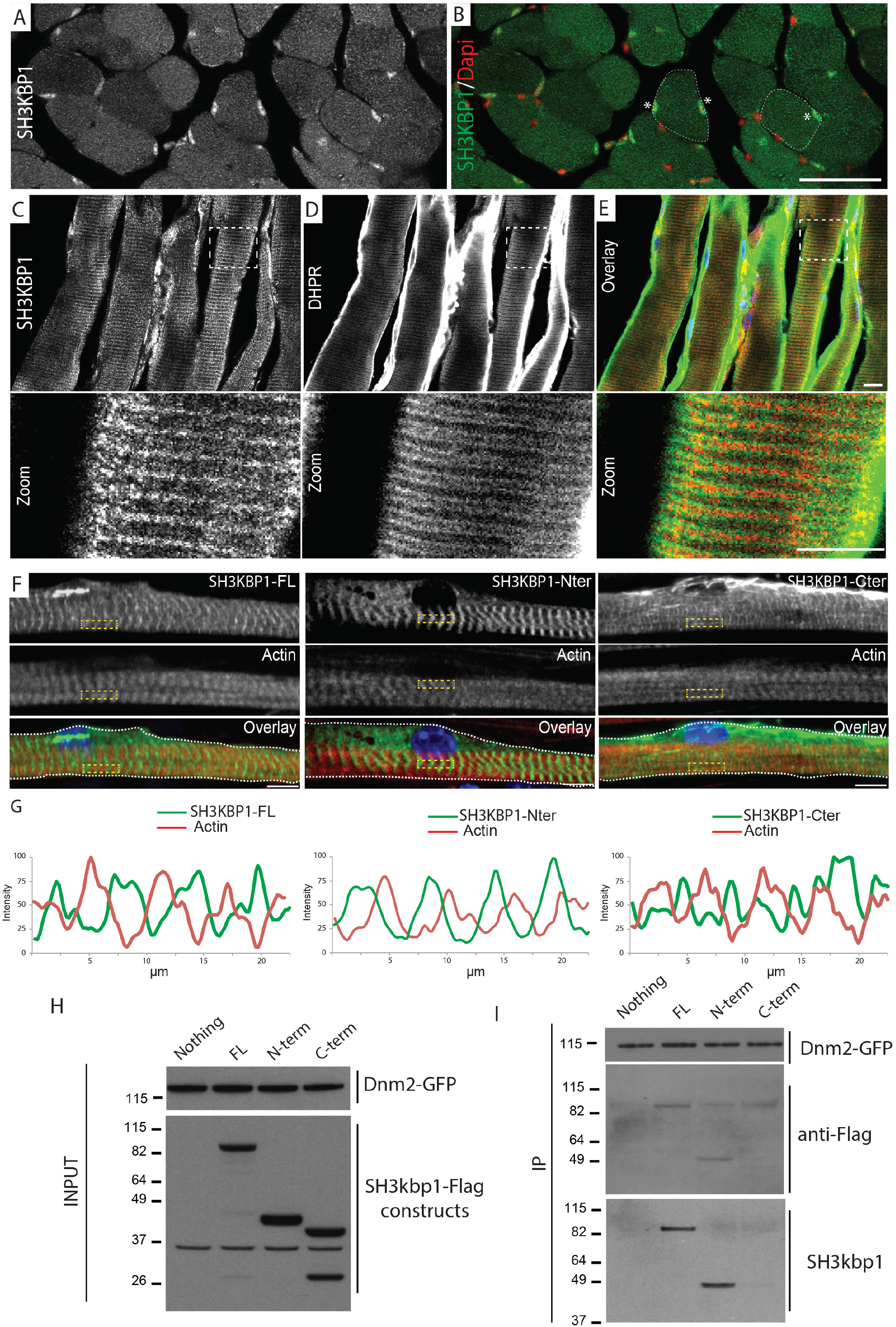
SH3KBP1 is localized near myonuclei and at I-band/Z-line zone in muscle fibers *in vivo* and interact with DNM2. (A-B) Representative images of *Tibialis Anterior* muscle transversals cross-section stained for SH3KBP1 (green) and myonuclei (red). Asterisks show myonuclei inside myofibers. Scale bars, 150 μm. (C-E) Representative images of *Tibialis Anterior* muscle longitudinal cross-section stained for SH3KBP1 (red) and DHPR1α, for DyHydroPyridine Receptor alpha (green). Scale bars, 150 μm. (F) Representative immunofluorescent staining of long term expression of SH3KBP1 constructs (listed in Fig. 4E) in green (GFP-constructs), Actin (red) and myonuclei (blue) in 10 days differentiated primary myofibers. Scale bars, 10 μm. (G) Line scan of the yellow boxes to visualize transversal organization of SH3KBP1 (green) and actin (red) staining. (H) Western blot with indicated antibodies (right) in C2C12 cells expressing Flag-SH3KBP1 constructs (top) and GFP-DNM2. (I) Representative western blot with indicated antibodies (right) of DNM2-GFP constructs immunoprecipitation using C2C12 expressing indicated constructs (top).

### SH3KBP1 is required in mature fibers for the maintenance of T-tubules

We next assessed the impact of *Sh3kbp1* depletion on internal cell architecture. To this end, we used our *in vitro* model assay on mature fibers using primary myoblasts (Fig. 6A). Myofibrillogenesis in *Sh3kbp1* depletion condition was normal after 10 days of differentiation, reflected by F-actin staining (Fig. 6B). In control condition, we also observed that DHPRα staining which labels T-tubule, forms transversal doublet bands alternatively with actin staining (Fig. 6A). Quantification of the staining indicated that in control conditions, using either siRNA or shRNA scramble, nearly 30% of formed myofibers exhibit a transversal DHPRα staining, reflecting formation of mature T-tubules (Fig. 6A & C). On the contrary, in *Sh3kbp1*-depleted myofibers, DHPRα staining is much more punctuated, indicating the absence of mature T-tubules aligned with sarcomere structures (Fig. 6B) and suggesting a role of SH3KBP1 in T-tubule formation. Indeed, in *Sh3kbp1*-myofibers, we observed a drop of myofibers with transversal staining of DHPRα to nearly 5% of myofibers, correlating with an increase in random DHPRα staining as dots along myofibers (Fig. 6C, Fig. S2E). To confirm these results we depleted *Sh3kbp1* from mature myofibers using shRNA electroporation technique directly in TA skeletal muscle of two-month-old mice (Fig. 6D). First, after 15 days of *Sh3kbp1* inhibition, an atrophic muscle fiber effect was observed in electroporated TA muscles fibers, reflected by a 35% decrease of the mean muscle fibers cross section area in *Sh3kbp1* fibers compared to controls (Fig. 6F), an effect that we previously observed *in vitro* (Fig. 2I). Moreover, T-tubule architecture appears perturbed in *Sh3kbp1* fibers, illustrated by an unequal DHPRα staining along fibers (Fig. 6D). Together, these results show that Sh3kbp1 contribute to T-tubule formation and maintenance in skeletal muscle fibers.

**Figure 6:**
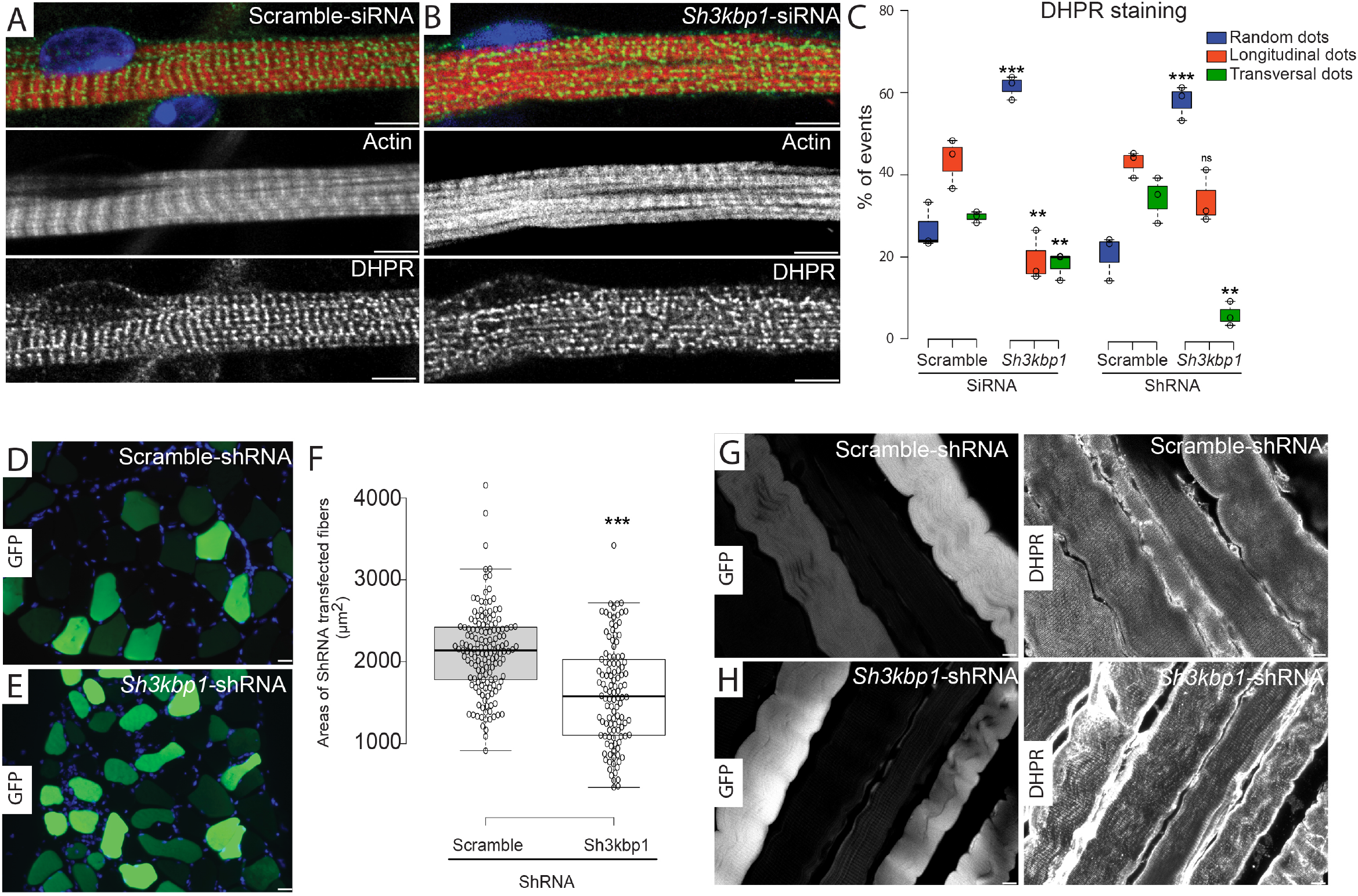
SH3KBP1 controls T-tubule organization during muscle fibers formation and in mature myofibers. (A-B) Representative immunofluorescent image of DHPR1-α (green), Actin (red) and myonuclei (blue) staining in 10 days cultured primary myotubes after scramble siRNA (A) or *Sh3kbp1* siRNA (B) transfection. Scale bars, 20 μm. (C) Quantification of DHPR-α staining aspects in mature myofibers treated with either scramble siRNA, a pool of 2 individual siRNAs targeting *Sh3kbp1* mRNA, scramble shRNA or an individual shRNA targeting *Sh3kbp1* mRNA. Data from three independent experiments were combined. Unpaired t-test, ***p < 0.001, **p < 0.01. Details for different categories are available in Fig. S2E. (D-E) Representative images of transversals (left) cross-section of *Tibialis Anterior* muscle electroporated with scramble or an individual shRNAs targeting *Sh3kbp1 and* stained for GFP (green) and myonuclei (blue). Scale bars, 50 μm (F) Quantification of muscle fibers electroporated with scramble or an individual shRNAs targeting *Sh3kbp1*. Center lines show the medians; box limits indicate the 25th and 75th percentiles as determined by R software; whiskers extend 1.5 times the interquartile range from the 25th and 75th percentiles, outliers are represented by dots. Unpaired t-test, ***p < 0.001. (G-H) Representative images of longitudinal cross-section of *Tibialis Anterior* muscle electroporated with scramble or an individual shRNAs targeting *Sh3kbp1 and* stained for GFP and DHPR-α. Scale bars, 20 μm.

### *Sh3kbp1* is up regulated in a murine model of AD-CNM and inhibits CNM phenotypes

Our data show that *Sh3kbp1* controls myonuclei dynamics and reticulum spatial organization and interacts with DNM2 which is mutated in dominant centronuclear myopathy characterized by myonuclear centralization (Romero & Bitoun, 2011). Heterozygous AD-CNM-Knock-In-*Dnm2*^R465W^ mouse model (KI-*Dnm2*^R465W/+^) progressively develop muscle atrophy, impairment of contractile properties, histopathological abnormalities including slight disorganization of T-tubules and reticulum, and elevated cytosolic calcium concentration (Durieux *et al*, 2010). We first analyzed *Sh3kbp1* mRNA expression in TA skeletal muscle during mice development and aging (Fig. 7A). To our surprise, *Sh3kbp1* mRNA was progressively reduced from more than 30 % in 8 months old mice compare to 2 months old mice (Fig. 7A), suggesting a progressive loss of SH3KBP1 pool during aging. More interestingly, we also showed that in this KI-*Dnm2*^R465W/+^ model, *Sh3kbp1* mRNA is significantly up-regulated during the first 4 months of mice development (Fig. 7A), suggesting a compensation mechanism, that can explain the limited ratio of atrophic fibers and absence of centralized myonuclei in this CNM model (Durieux *et al*, 2010). To address the role of SH3KBP1 in long-term muscle homeostasis in both wild type and KI-*Dnm2*^R465W^ mice, we investigated *in vivo* depletion of Sh3kbp1 protein. Down-regulation of *Sh3kbp1* was achieved using intramuscular TA muscle injections of an AAV cognate vector expressing shRNA targeting *Sh3kbp1* mRNA (AAV-shSh3kbp1) either in wild type or KI-*Dnm2*^R465W^ mice at 5 weeks of age, an age where muscle mass is nearly fully developed. This allowed addressing the role of *Sh3kbp1* specifically in adult skeletal muscle during the first three months of mice development and specifically in a period where we observed *Sh3kbp1* mRNA up-regulation in KI-*Dnm2*^R465W^ model (Fig. 7A). In wild type mice, *Sh3kbp1* mRNA level was decreased by 2.7 fold compared to PBS-injected muscles (Fig. S3A, WT). In KI-*Dnm2*^R465W^ mice, *Sh3kbp1* mRNA levels showed a 5.4 fold decrease compared to PBS-injected TA-muscles (Fig. S3A, WT, KI-*Dnm2*^R465W^). No significant change of body weight was observed between AAV-shSh3kbp1-injected and PBS-injected conditions in both genotypes (Fig. S3B). However, a significant decrease in absolute force (g) developed by TA muscles specifically in KI-*Dnm2*^R465W^ mice depleted for *Sh3kbp1* was observed (Fig. 7D) suggesting a specific impact of *Sh3kbp1* down-regulation in the KI-*Dnm2*^R465W^ mice model. Interestingly, we observed a significant decrease of TA muscle mass of about 20%, when *Sh3kbp1* is depleted in both WT and KI-*Dnm2*^R465W^ mice (Fig. 7E). Cross-sectional areas of TA-muscle fibers were determined using transverse sections stained with laminin antibodies to define muscle fibers border (Fig. 7B-C). Global decrease in median myofibers area was observed in AAV-shSh3kbp1 injected TA-muscles compared to control muscles in WT (−30%) and KI-*Dnm2*^R465W^ (−20%) mice (Fig. 7F). Analysis of muscle fibers area repartition showed a significant increase in atrophic fibers in AAV-shSh3kbp1 injected muscles (Fig. S3C). Additionally, we observed a significant reduction of about 25% in the total number of fibers specifically in KI-*Dnm2*^R465W^ model depleted for *Sh3kbp1* (Fig. S3D). DNM2-CNM patients exhibit a predominance of type 1 muscle fibers (Romero & Bitoun, 2011). Thus, we investigated the ratio of slow fiber types in AAV-shSh3kbp1 injected TA-muscles, reflected by the expression of myosin heavy chain type 1 (Fig. 7C-D & G). In wild type conditions, slow fiber type accounted for 5 % of total muscle fibers. We observed that in AAV-shSh3kbp1 injected muscles, this ratio was slightly increased to reach 6.4 % of total muscle fibers, which is the same as what was observed in KI-*Dnm2*^R465W^ mice (Fig. 7G). However, when *Sh3kbp1* is depleted in KI-*Dnm2*^R465W^ muscle, this ratio increases drastically to reach 12% of total fibers (Fig. 7G). In the Knock-in mouse model, DNM2 mutation leads to autophagy impairment (Rabai *et al*, 2019; Durieux *et al*, 2010). During autophagosome formation, LC3-I (Microtubule-associated protein 1A/1B-light chain 3) is converted to LC3-II by conjugation to phosphatidylethanolamine lipid. To assess the impact of *Sh3kbp1* depletion on autophagy, LC3-II positive fibers was determined in Sh3kbp1 depleted fibers in both WT and KI-Dnm2R465W mice. We observed an increase in number of LC3-II positive fibers in both genotypes but more pronounced in KI-*Dnm2*^R465W^ mice (Fig. 7H). Our *in vitro* data show that *Sh3kbp1* depletion increase fusion and alter myonuclei spreading (Fig. 1-3). We thus investigate the impact on the number of myonuclei and the ratio of internalized myonuclei by fibers in *Sh3kbp1* depleted myofibers but did not find any evidence for alteration compare to control conditions (Fig. S3E-F). Altogether, our data suggest that *Sh3kbp1* mRNA increase observed in AD-CNM model could be an attempt of compensatory mechanism as normalization of *Sh3kbp1* expression intensified muscle phenotype.

**Figure 7:**
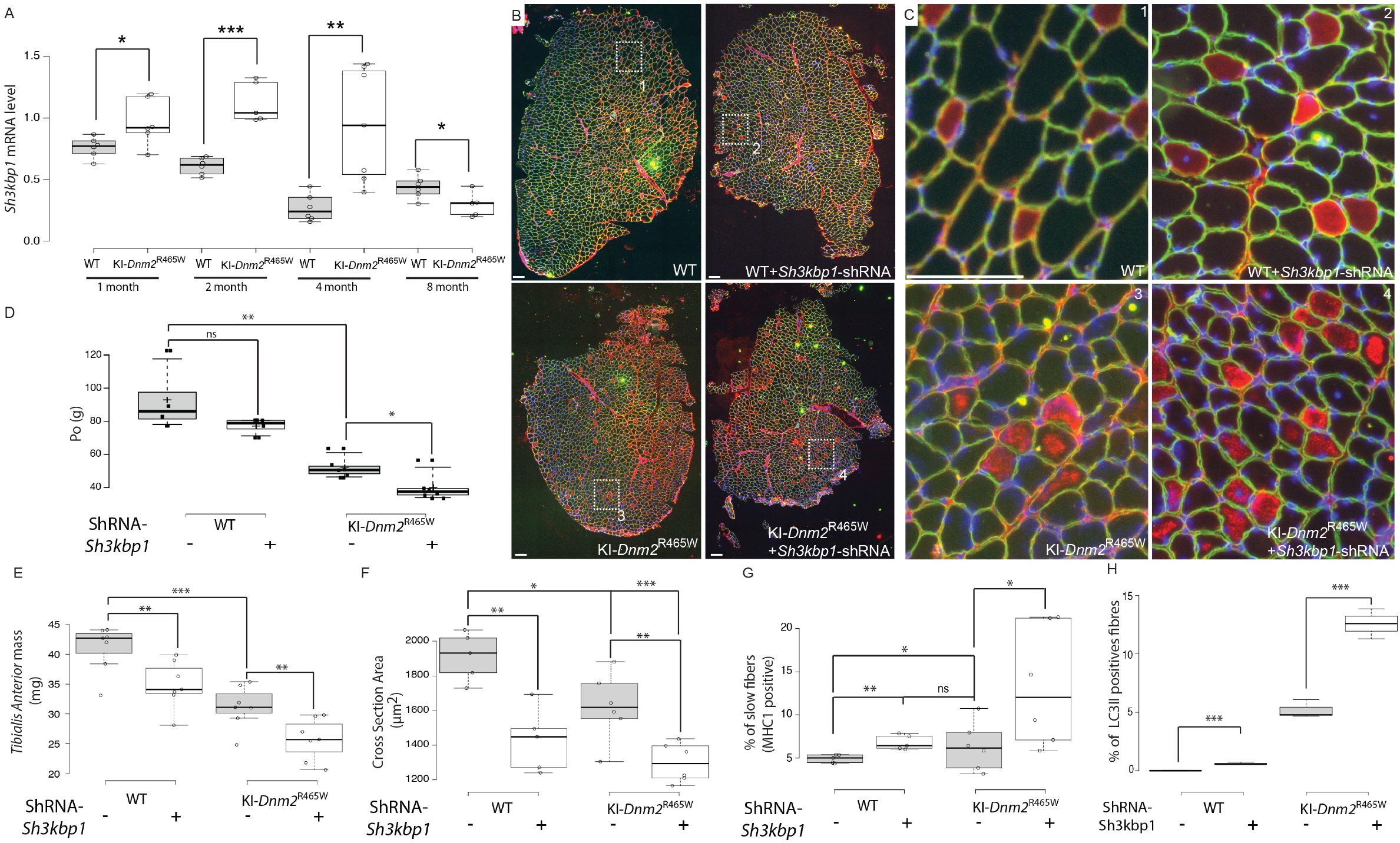
*sh3kbp1* silencing worsened CNM phenotype in KI-DNM2^R465W^ mice model. (A) qRT-PCR analysis of *Sh3kbp1* gene expression relative to *Nat10* gene *Tibialis Anterior* muscles from WT or KI-*Dnm2*^R465W^ mice at 1, 2, 4 and 8 months of age. Non-parametric one-way Anova used for statistical significance ***p < 0.001, **p < 0.01, *p < 0.05. (B-C) Representative images of *Tibialis Anterior* muscle cross-section from WT or KI-*Dnm2*^R465W^ mice at the age of 4 months, injected with either PBS or AAV cognate vector expressing shRNA targeting *SH3KBP1* mRNA (Sh3kbp1-shRNA) for 3 months and stained for Dapi (Blue), Laminin (green) and MyHC (red). Scale Bars = 150 μm. (D-H) Quantification of Absolute force (P0) (D), *Tibialis Anterior* muscle mass (E), mean cross-sectioned myofibers areas (F), percentage of slow fibers (MHC1 positives) (G) and percentage of LC3II-positive fibers (H) of *Tibialis Anterior* muscles from WT or KI-*Dnm2*^R465W^ mice at the age of 4 months, injected with either PBS or AAV cognate vector expressing shRNA targeting *SH3KBP1* mRNA (Sh3kbp1-shRNA) for 3 months. Data from at least 4 mice in each condition were combined. Unpaired t-test, ***p < 0.001, **p < 0.01, *p < 0.05. Center lines show the medians; box limits indicate the 25th and 75th percentiles as determined by R software; whiskers extend 1.5 times the interquartile range from the 25th and 75th percentiles, outliers are represented by dots.

## Discussion

Myonuclei movement’s during muscle cell formation have been related to the activity of numerous microtubule binding proteins such as Motors and Maps (Gache *et al*, 2017). In immature myotubes, microtubules are organized into antiparallel arrays between adjacent myonuclei and contribute to the polarization/elongation of myotubes, alignment of myonuclei and ultimately organization of myosin filaments during sarcomere formation (Pizon *et al*, 2005; Metzger *et al*, 2012; Wang *et al*, 2013). This new orientation/architecture of the microtubule network directly influences myoblasts/myotubes fusiform shape, as microtubule network forces are applied at the extremity of polarized myotubes and also in-between myonuclei (Metzger *et al*, 2012); it also contributes to myonuclei spreading along myotubes length (Tassin *et al*, 1985). This precise myonuclei organization in muscle fibers gives rise to the formation of so-called “MyoNuclear Domains” (MNDs) wherein each myonucleus is responsible for gene expression in its surrounding cytoplasm and guaranties functional integrity of muscles (Qaisar, 2012; Liu *et al*, 2009). However, regulation of processes involved in myonuclei positioning in developing muscle fibers, not directly linked to the microtubule network, are still unknown.

The present study demonstrates that a protein related to the endoplasmic reticulum (ER) network controls both myotubes elongation and myonuclei localization/spreading in developing myofibers, two processes that are closely linked as they both depends on microtubule network organization. Early steps of myonuclei positioning in myotubes depend on an interplay between microtubules, microtubules-associated-proteins such as MAP4, MAP7 and motors proteins such as Dynein and Kif5B (Gimpel *et al*, 2017; Mogessie *et al*, 2015; Metzger *et al*, 2012; Cadot *et al*, 2012; Wang *et al*, 2013; Wilson & Holzbaur, 2014a). This process also depends on myonuclei membrane associated components such as Nesprin-1 or AKAP9 (Gimpel *et al*, 2017; Wilson & Holzbaur, 2014b; Doñate Puertas *et al*, 2018). Here, we show that SH3KBP1 depletion leads to longer and thinner myotubes without significant alteration of the microtubules network compared to control (Fig. 1-3 & data not shown). We find that the C-terminus part of SH3KBP1 is associated with ERP72-positive ER probably through its binding to Calnexin (Fig. 4E-G). This association scaffolds ERP72-associated ER specifically at myonuclei vicinity, without affecting Golgi apparatus integrity (Fig. 4A-D). Recently, ER has been implicated in governing both microtubule alignment and cytoplasmic streaming (Kimura *et al*, 2017). Kimura *et al* propose that local cytoplasmic flow generated along microtubules is transmitted to neighboring regions through the ER and in turn, aligns microtubules and self-organizes the collective cytoplasmic flow. SH3KBP1, by shaping and maintaining ER clustering during myofibers development could contribute to prevent microtubules network anchoring at the nucleus membrane and thus local microtubule network organization. In this view, SH3KBP1, during myoblasts fusion process could disorganize parallel microtubules organization and thus decrease forces applied on myotubes tips extremity. This phenomenon could lead to the control of myotubes length, but also of both myonuclei motility and velocity (Fig. 2).

SH3KBP1 staining is diffuse in dividing cells, while it progressively accumulates at the vicinity of myonuclei during differentiation process (Fig. 4). A few proteins have been shown to form a flexible perinuclear shield that can protect myonuclei from extrinsic forces (Wang *et al*, 2015; Ghasemizadeh *et al*, 2019). To note, recent data show that disruption of one component of this perinuclear shield, MACF1 protein, increases myonuclei velocity (Ghasemizadeh *et al*, 2019). Consequently, SH3KBP1 probably belongs to the group of proteins that contribute to the stability of this perinuclear shield through the maintenance of ER at the vicinity of myonuclei. In accordance with this preferential localization at the proximity of myonuclei, SH3KBP1 staining in transversal *Tibialis Anterior* muscle section is very intense at myonucleus site whereas nuclei outside the fiber remains negative (Fig. 5A-B). In human muscle biopsies, this localization around myonuclei is maintained even in centralized nuclei from CNM patient (Fig. S3G). The three SH3 domains of SH3KBP1 have been shown to cluster multiple proteins and protein complexes that can also contributes to the stability of those interactions. Havrylov *et al* identified using mass spectrometry, few microtubule-binding-proteins such as MAP7 and MAP4 that can potentially interact with SH3 domains of SH3KBP1 and have already been shown to control myonuclei positioning in myotubes (Metzger *et al*, 2012; Mogessie *et al*, 2015; Havrylov *et al*, 2009). However, we failed to confirm the interaction of SH3KBP1 with either MAP7 or MAP4 in muscle cells (data not shown). Alternatively, SH3KBP1 also interacts with dynamin-2 (DNM2) and has been shown to participate in dynamics instability of microtubules (Tanabe & Takei, 2009) and microtubule nucleation (Thompson *et al*, 2004). The failure of recruiting DNM2 at the right place during myotubes formation after SH3KBP1 depletion could also contribute to the aberrant myonuclei spreading (Fig. 1). Interestingly, in CNM-KI-*Dnm2*^R465W^ mouse model, Fongy *et al* show that myonuclei move and spread properly in heterozygous myotubes but hypothesize a defect in nuclear anchoring at the periphery (Fongy *et al*, 2019). Several studies pointed the importance of cytoskeleton, including MAPs, microtubules and intermediate filaments, in the nuclear anchorage in mature muscle (Ghasemizadeh *et al*, 2019; Roman *et al*, 2017). SH3KBP1 dependent ER-scaffolding could participate in myonuclei anchoring at the periphery of myofibers and thus in the recruitment and stabilization of a network of proteins at the vicinity of myonuclei. This hypothesis is supported by our data showing an increase in the percentage of the time in motion of myonuclei, in the absence of SH3KBP1 (Fig. 2D), correlated with the strong staining of myonuclei in mouse and human models (Fig. 5A & S3G).

In mature myofibers, SH3KBP1 depletion leads to more aggregated myonuclei phenotype, suggesting that a failure in the early phases of myonuclear positioning is difficult to compensate (Fig.2G). Interestingly, *sh3kbp1* depletion in long-term primary myofibers culture induces a reduction in myofibers width (Fig. 2I). These results are confirmed *in vivo* as *sh3kbp1* down regulation in *Tibialis Anterior* muscle reduce from more than 30% the average cross section areas of myofibers (Fig. 6E-F & 7F). Moreover, SH3KBP1-depleted mature myofibers show disorganized perinuclear endoplasmic reticulum (ER) (Fig 4). Of interest, ER is one initiation site for autophagic process and ER selective autophagy (called ER-phagy or reticulophagy) has been described to control ER shape and dynamics through ER-phagy receptors that address ER portions to the autophagosomes (Grumati *et al*, 2018). SH3KBP1 depletion increases the number of LC3II positives myofibers in *Tibialis Anterior* muscle (Fig. 7H), suggesting a possible increase of autophagosomes number. Additionally, SH3KBP1 interacts with dynamin-2 (Fig. 5-G), which is also involved in the autophagic lysosome reformation (Schulze *et al*, 2013). This autophagy-dependent modulation of muscle homeostasis could first explain the decrease of Cross Section Area and myofibers width that we observed and ultimately the reduction of muscle myofibers (Fig. 2I, 7E-F & S3C). Interestingly, autophagy genes have been involved in muscle myonuclei positioning during Drosophila metamorphosis (Fujita *et al*, 2017). Whether this process is involved in SH3KBP1-dependent myonuclei positioning will be the subject of further investigations.

SH3KBP1 depletion shows that T-tubule formation is dramatically impaired both *in vitro* and *in vivo* (Fig. 6). Myofibrils provide the contractile force under the control of the ‘excitation-contraction coupling’ system that includes two membranous organelles: the sarcoplasmic reticulum (SR) and Transverse (T)-tubules (Al-Qusairi & Laporte, 2011). These two-membrane systems are structurally associated to form the triads of skeletal muscle cells. SR is a complex network of specialized smooth endoplasmic reticulum, essential to transmit the electrical impulse as well as in the storage of calcium ions. SR is built from the formation and maturation of two distinct and functionally related domains: the longitudinal SR and the junctional SR that together wrap the contractile apparatus. T-tubule network is continuous with the muscle cell plasma membrane (PM) and begins from the invagination of PM in a repeated pattern at each sarcomere (Barone *et al*, 2015). These two-membrane systems are structurally associated to maintain their typical organization in muscle cells. SH3KBP1 depletion seems to alter more specifically junctional SR rather than longitudinal SR as it specifically alters transversal organization of T-tubules (Fig. 6A-C). One hypothesis, recently suggested by Quon *et al*, is that non-vesicular lipid transport and lipid biosynthesis could intersect at ER-PM membrane contact sites and would serve as a nexus, coordinating requirements in the PM for lipids with their production in the ER (Quon *et al*, 2018).

Autosomal dominant CNM is caused by heterozygous mutations in the *DNM2* gene, which encodes the Dynamin 2 (DNM2) GTPase enzyme (Romero, 2010). *DNM2*-related autosomal dominant (AD)-CNM was initially characterized as a slowly progressing muscle weakness affecting distal muscles with onset in early adulthood. DNM2-R465W missense mutation represents the most frequent mutation in humans and a knock-in (KI) mouse model expressing this mutation has been generated that develops a progressive muscle weakness (Durieux *et al*, 2010). Expressing DNM2-R465W mutation in mice leads to contractile impairment that precedes muscle atrophy and structural disorganization that mainly affects both mitochondria and endo/sarcoplasmic reticulum. Interestingly, CNM-KI-*Dnm2*^R465W^ mouse model exhibits twice more Sh3kbp1 mRNA amount in the first 4 months than control mice before normalization after 8 months (Fig. 7A). Of note, this increase is concomitant with transient transcriptional activation of both ubiquitin–proteasome and autophagy pathways at 2 months of age in the TA muscle of the KI-Dnm2 mice (Durieux *et al*, 2010). One can hypothesize that this elevated amount of *sh3kbp1* is one of the factors that limits activation of autophagy pathway activation and thus slows-down CNM associated phenotype. In accordance with this hypothesis, we find that *sh3kbp1* depletion increase CNM phenotype (Fig. 7D-H). In conclusion, increased amount of SH3KBP1 could delay the CNM phenotype development by stabilizing Triads and the braking of the autophagy response.

Finally, our data show that *Sh3kbp1* depletion increases fusion events in transfected primary myoblasts and stable cell line. *In vivo,* when *Sh3kbp1* is depleted from mature myofibers in a period of time characterized by minimal myonuclei accretion in muscle fibers, we observe a tendency to slightly increase the percentage of myonuclei per myofibers in both WT and KI-*Dnm2*^R465W^ conditions (Fig. S3E). These data suggest that SH3KBP1 could contribute to fusion efficiency by a still unknown mechanism. The role of ER on myoblast fusion is poorly documented and its effect seems to be more indirectly linked to pathways related to physiologic ER stress signaling and SARC (stress-activated response to Ca^2+^) body formation more than a direct impact on myoblast fusion (Bohnert *et al*, 2017; Nakanishi *et al*, 2007; 2015). Alternatively, our *in vitro* data suggest that the global microtubule network organization is not changed but that dynamics of organelles related to microtubules pathways are improved, reflected by the increase of myonuclei dynamics in the absence of SH3KBP1 (Fig. 2A-F). There is also no evidence of a clear specific role of microtubules network on the fusion potential, however, the modification of microtubule dynamics/orientation by different MAPs are known to alter fusion potential (Mogessie *et al*, 2015; Straube & Merdes, 2007; Cadot *et al*, 2012). Fusion processes require remodeling of both membranes and actin cytoskeleton polymerization at the site where two membrane will fuse (Sampath *et al*, 2018). One can hypothesize that SH3KBP1 could contribute to the fusion process through the control of membrane remodeling on specifics PM-ER sites. Incidentally, we noticed that in absence of SH3KBP1, myotubes are longer, increasing consequently the fusiform shape of cells (Fig. 1D). As fusion mainly occurs at the tips of myotubes (Cadot *et al*, 2015; 2012), this increase of myotubes polarization could contribute to accumulate proteins involved in fusion specifically at the tips of myotubes and consequently favor fusion capacity.

Altogether these data are in agreement with an involvement of *Sh3kbp1* in muscle fibers formation and maintenance. In the present study, we show that the adaptor protein SH3KBP1, through its N-terminus part, is able to scaffold ER *via* an interaction with Calnexin. SH3KBP1 also governs myonuclei motion and spatial organization as well as myoblast fusion. Additionally, we observed that SH3KBP1 interacts with DNM2 and organizes T-tubule formation in skeletal muscle; moreover, its down-regulation contributes to an increase of the CNM-like phenotype in the most frequent mutation model (KI-*Dnm2*^R465W^) of autosomal dominant centronuclear myopathy. Altogether, our data suggest that SH3KBP1 increase observed in AD-CNM model could be an attempt of compensatory mechanism in CNM-KI-*Dnm2*^R465W^ model and could be used as a preventing factor in the development of CNM phenotype.

---

**Supplementary figure 1:**
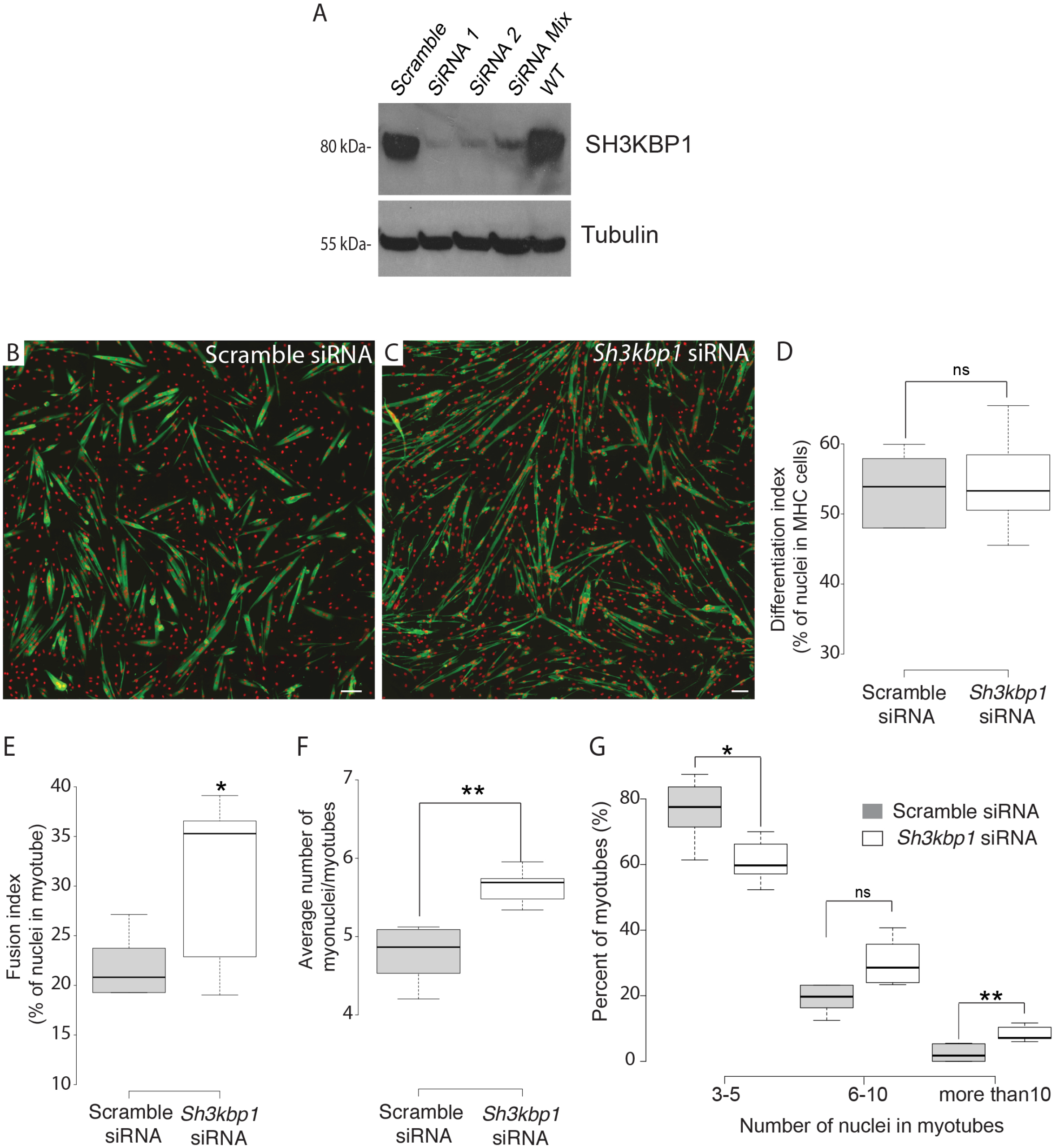
SH3KBP1 affects myoblast fusion. (A) Representative western blot analysis of SH3KBP1 protein expression in total protein extracts from differentiating primary myotubes treated with scramble or siRNA (#1 & #2) or a pool of siRNA targeting *SH3KBP1* mRNA. Loading control is tubulin. (B-C) Representative immunofluorescent images of 3 days of primary differentiated myotubes stained for myosin heavy chain (green) and myonuclei (red) and treated with scramble (B) or with a pool of 2 individual *Sh3kbp1* siRNAs. Scale Bar: 50μm. (D-G) Quantification of differentiation index (percentage of nuclei in Myosin Heavy Chain positives cells) (D), Fusion index (percentage of nuclei inside myotubes) (E), Average number of nuclei by myotubes (F) and distribution of myotubes classified depending on the number of nuclei in myotubes (G). Data from three independent experiments were combined. Unpaired t-test, **p < 0.01, *p < 0.05. Center lines show the medians; box limits indicate the 25th and 75th percentiles as determined by R software; whiskers extend 1.5 times the interquartile range from the 25th and 75th percentiles, outliers are represented by dots.

**Supplementary figure 2:**
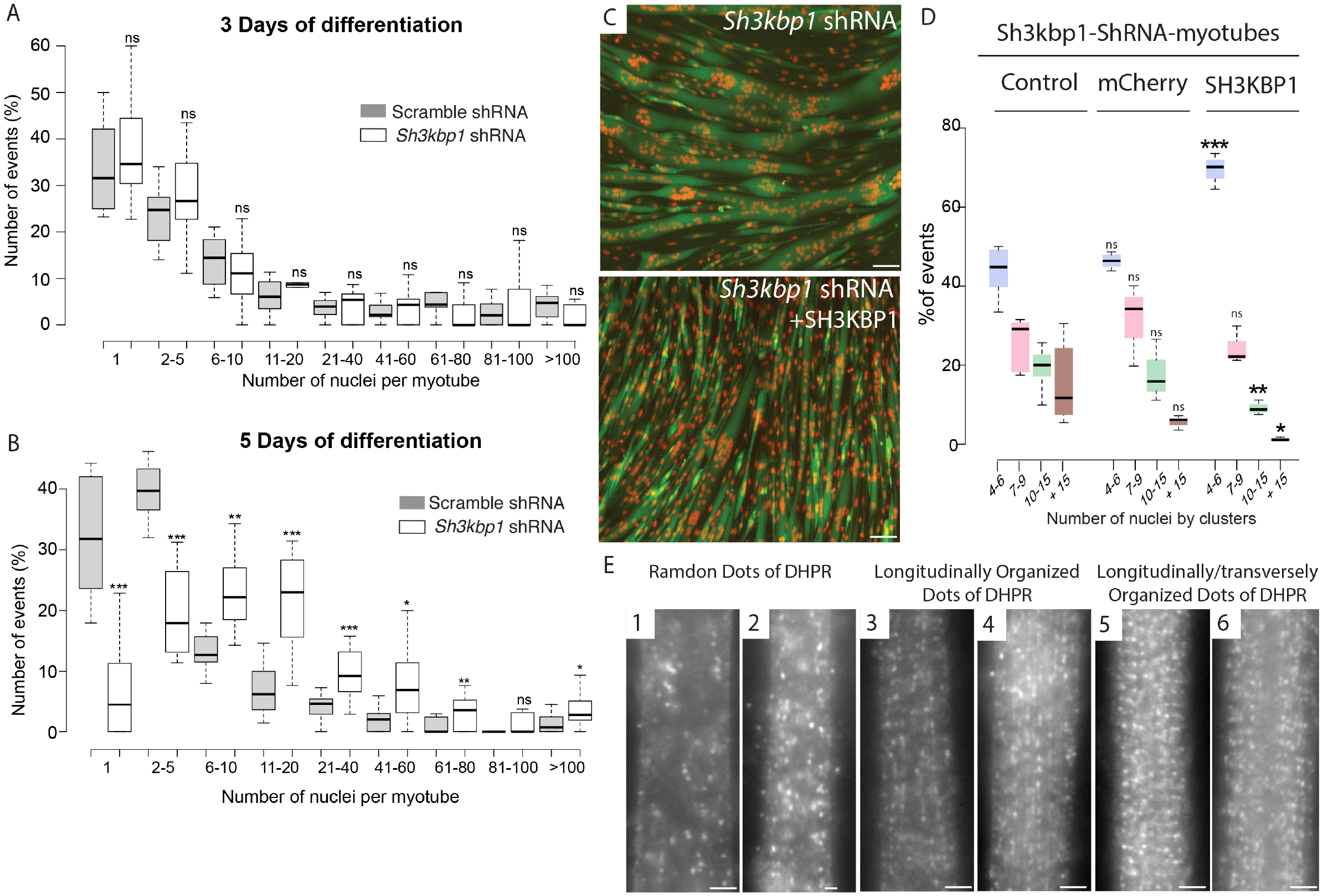
SH3KBP1 affects myoblast fusion and localization of myonuclei in C2C12 cells. (A-B) Distribution of 3 days differentiated C2C12 myotubes formed from stable cell line expressing either scramble or *Sh3kbp1* shRNAs and classified depending on nuclei content inside myotubes after (A) 3 days or (B) 5 days of differentiation. (C) Representatives immunofluorescent staining of myosin heavy chain (green) and myonuclei (red) in stable cell line expressing *Sh3kbp1* shRNA (top) or co-transfected with full-length SH3KBP1 (bottom) after 5 days of differentiation. Scale Bar: 50μm. (D) Quantification of the percentage of nuclei by clusters in stable cell line expressing *Sh3kbp1* shRNA and co-transfected with mCherry or full-length SH3KBP1 plasmid. Data from three independent experiments were combined. Unpaired t-test, ***p < 0.001, **p < 0.01, *p < 0.05. Center lines show the medians; box limits indicate the 25th and 75th percentiles as determined by R software; whiskers extend 1.5 times the interquartile range from the 25th and 75th percentiles, outliers are represented by dots. (E) Representatives immunofluorescent categories of staining for DHPR-alpha in 10 days differentiated myofibers. Scale Bar: 2μm.

**Supplementary figure 3:**
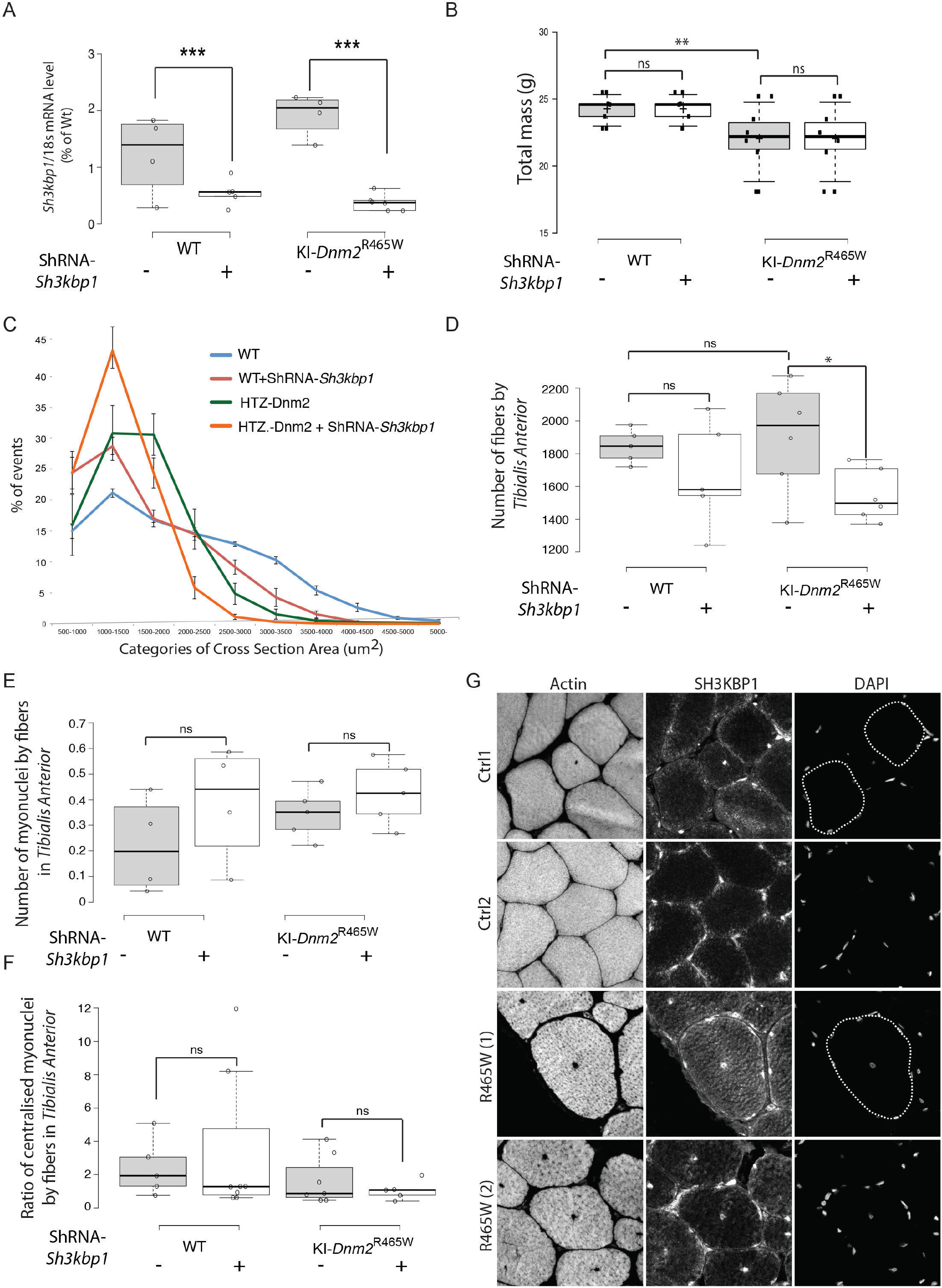
*sh3kbp1* silencing worsened CNM phenotype in KI-DNM2R465W mice model. (A) qRT-PCR analysis of *Sh3kbp1* gene expression relative to Nat10 gene in 4 months *Tibialis Anterior* muscles from WT or KI-*Dnm2*^R465W^ mice injected with either PBS or AAV cognate vector expressing shRNA targeting *SH3KBP1* mRNA (Sh3kbp1-shRNA). ***p < 0.001. (B-F) Quantification of total body mass (B), distribution of cross-sectioned myofibers areas (C), number of fibers by muscles (D), number of myonuclei by fiber (E) and number of internalized myonuclei (F) in *Tibialis Anterior* muscles from WT or KI-*Dnm2*^R465W^ mice at the age of 4 months, injected with either PBS or AAV cognate vector expressing shRNA targeting *SH3KBP1* mRNA (AAV-SH3KBP1) for 3 months. Data from at least 4 mice in each condition were combined. Unpaired t-test, ***p < 0.001, **p < 0.01, *p < 0.05. Center lines show the medians; box limits indicate the 25th and 75th percentiles as determined by R software; whiskers extend 1.5 times the interquartile range from the 25th and 75th percentiles, outliers are represented by dots. (G) Immunofluorescent staining of SH3KBP1, Actin and nuclei in two control human patient and in two patients expressing DNM2-R465W point mutation.

**Supplementary video 1 & 2:** Timelapse experiments of primary myoblasts co-transfected with Lamin-Chromobody®-RFP plasmids and with either Scramble-siRNA (Supplementary video-1) or *Sh3kbp1-*siRNA (Supplementary video-2) and induce to form myotubes during 5 days after starting differentiation process. Primary myotubes were recorded every 20 minutes for a period of time of 16 hours.

## Materials and methods

### Cell culture

Primary myoblasts were collected from wild type C57BL6 mice as described before (Falcone *et al*, 2014; Pimentel *et al*, 2017). Briefly, Hindlimb muscles from 6 days pups were extracted and digested with collagenase (Sigma, C9263-1G) and dispase (Roche, 04942078001). After a pre-plating step to discard contaminant cells such as fibroblasts, myoblasts were cultured on matrigel coated-dish (Corning, 356231) and induced to differentiate in myotubes for 2-3 days in differentiation media (DM: IMDM (Gibco, 21980-032) + 2% of horse serum (Gibco, 16050-122) + 1% penicillin-streptomycin (Gibco, 15140-122)). Myotubes were then covered by a concentrated layer of matrigel and maintained for up to 10 days in long differentiation culture medium (LDM: IMDM (Gibco, 21980-032) + 2% of horse serum (Gibco, 16050-122) + 0.1% Agrin + 1% penicillin-streptomycin (Gibco, 15140-122)) until the formation of mature and contracting myofibers. LDM was changed every two days.

Mouse myoblast C2C12 cells were cultured in Dulbecco’s modified Eagle’s medium (DMEM (Gibco, 41966029) + 15% fetal bovine serum (FBS) (Gibco, 10270-106) + 1% penicillin-streptomycin (Gibco, 15140-122))) and were plated on 0.1% matrigel-coated dishes for 1-2 days before differentiation. Differentiation was induced by switching to differentiation media (DMEM + 1% horse serum).

### Cell transfection

For C2C12 cells, 3 different siRNAs Silencer per gene were transfected using Lipofectamine 2000 (ThermoFisher Scientifics, 11668-019) at the final concentration of 10 nM, following manufacturer instructions, 2 days before differentiation. For shRNA cDNA (Geneocopia) transfection, Lipofectamine 2000 (ThermoFisher Scientifics, 11668-019) was used following manufacturer instructions.

For primary cells, siRNA were transfected using Lipofectamine 2000 (ThermoFisher Scientifics, 11668-019) at the final concentration of 2 nM. shRNA (Geneocopia), Eb1 or RFP-Lamin-chromobody (Chromotek) cDNA were transfected using Lipofectamine 3000 (ThermoFisher Scientifics, L3000-008). For the list of siRNA, shRNA and DNA constructs, refer to supplementary table 1.

### Protein sample preparation

For primary cultured cells or C2C12 cell lines, cells were harvested, using Trypsin for 5min at 37°C and centrifuged at 1500RPM for 5min at 4°C. Cell pellets were diluted and incubated in the optimal volume of RIPA lysis buffer containing phosphatases inhibitors (Sigma, P5726-5mL) and proteases inhibitors (Sigma, P8340) for 10min at 4°C. Following a sonication and a centrifugation at 12000RPM for 10min at 4°C, protein samples were collected for further uses. The concentration of proteins was determined using BCA protein assay kit (Thermo Fisher Scientifics, 23225) as described by the manufacturer.

### Western blot

To carry out western blots, the same amount of sample were loaded in 6% acrylamide gels and were migrated at 130V for 10min followed by 160V for 90min. iBlot 2 mini slacks (Thermo Fisher Scientifics, IB23002) semi-dry system was used to transfer the proteins to nitrocellulose membranes. Membranes were then saturated in 5% milk in TBS for 1h at room temperature (RT) and were incubated in primary antibodies in 5% milk in TBS over night at 4°C. Following washes by 0.1% Tween-20-1X TBS, the membranes were incubated in HRP conjugated secondary antibodies in 5% milk in TBST for 1h at room temperature (RT). Following washes by 0.1% Tween-20-1X TBS the detection of the target proteins was carried out using Super Signal West Femto (Thermo Fisher Scientifics, 34095) and ChemiDoc imaging system (BioRad).

### Antibodies

Cells were fixed in 4%PFA in PBS for 20min at 37°C followed by washes with PBS and permeabilization with 0.5% Triton-X100 in PBS for 5min at RT. Following washes with PBS, cells were saturated with 1% BSA in PBS for 30min at 37°C and incubated in primary antibodies over night at 4°C. Following washes with 0.05% Triton-X100 in PBS, cells were incubated in secondary antibodies or dyes for 2hrs at RT followed by washes with 0.05% Triton-X100 in PBS and a last wash in PBS. Cultured myofibers were imaged using either Z1-AxioObserver (Zeiss) or confocal SP5 microscope (Leica). For the list of antibodies and dilution, refer to supplementary table 2.

### Video-Microscopy

Time-lapse images were acquired using Z1-AxioObserver (Zeiss) with intervals of 20 minutes. Final videos were analyzed using Metamorph (Zeiss) and SkyPad plugin as described before (Cadot *et al*, 2014).

### Adeno-Associated Virus production and *in vivo* transduction

A cassette containing the small hairpin (sh) RNA under the control of H1 RNA polymerase III promoter was inserted in a pSMD2 expression plasmid. AAV vectors (serotype 1) were produced in HEK293 cells after transfection of the pSMD2-shRNA plasmid, the pXX6 plasmid coding for viral helper genes essential for AAV production and the pRepCap plasmid (p0001) coding for AAV1 capsid as described previously (Riviere et al, 2006). Viral particles were purified on iodixanol gradients and concentrated on Amicon Ultra-15 100K columns (Merck-Millipore). The concentration of viral genomes (vg/ml) was determined by quantitative real-time PCR on a LightCycler480 (Roche diagnostic, France) by using TaqMan probe. A control pSMD2 plasmid was tenfold serially diluted (from 10^7^ to 10^1^ copies) and used as a control to establish the standard curve for absolute quantification. Male wild type and heterozygous KI-*Dnm2*^R465W^ mice were injected under isoflurane anesthesia. Two intramuscular injections of 30 μl within 24h interval were performed using 29G needle in TA muscles corresponding to 10^11^ viral genomes per muscle. All of the experiments and procedures were conducted in accordance with the guidelines of the local animal ethics committee of the University Claude Bernard – Lyon 1 and in accordance with French and European legislation on animal experimentation and approved by the ethics committee CECCAPP and the French ministry of research.

### Muscle contractile properties

The isometric contractile properties of TA muscles were studied *in situ* on mice anesthetized with 60 mg/kg pentobarbital. The distal tendon of the TA muscle was attached to a lever arm of a servomotor system (305B Dual-Mode Lever, Aurora Scientific). The sciatic nerve was stimulated by a bipolar silver electrode using a supramaximal (10 V) square wave pulse of 0.1 ms duration. Absolute maximal isometric tetanic force was measured during isometric contractions in response to electrical stimulation (frequency of 25–150 Hz; train of stimulation of 500 ms). All isometric contraction measurements were made at optimal muscle length. Force are expressed in grams (1 gram = 9.8 mNewton). Mice were sacrificed by cervical dislocation and TA muscles were weighted. Specific maximal force was calculated by dividing absolute force by muscle weight.

### RNA extraction

After the addition of Trizol (Sigma, T9424-200mL) on each sample, lysing matrix D and fast prep system (MPbio, 6913-100) were used for sample digestion and pre-RNA extraction. In order to extract RNA, samples were incubated in chloroform for 5min at RT, centrifuged for 15min at 12000 rcf at 4°C and incubated in the tubes containing isopropanol (precipitatation of RNA) for 10min at RT. following a centrifuge of samples for 15min at 12000rcf at 4°C, samples were washed 2 times with 70% ethanol and the final RNA pellets were diluted in ultra-pure RNase free water (Invitrogen, 10977-035). RNA concentration was calculated using Nanodrop (ThermoFisher Scientifics).

### RT-q-PCR on cells

Goscript Reverse Transcriptase System (Promega, A5001) was used, as described by the manufacturer to produce the cDNA. Fast Start Universal SYBR Green Master (Rox)(Roche, 04913914001) and CFX Connect™ Real-Time PCR Detection System (BioRad) were used to carry out the quantitative PCR using the following primer sets. The CT of target genes were normalized on 3 control genes. For the list of primers used, refer to supplementary table 1.

### RT-q-PCR on muscle samples

50 longitudinal sections (12 μm) of TA muscles were cut and used for RNA isolation and RT-qPCR. Total RNA was extracted from muscle by using NucleoSpin (Macherey-Nagel). RNA (200 ng) was reverse transcribed using Reverse Transcription Core Kit (Eurogentec). Real-time PCR was performed in a 20 μL final volume using the Takyon No Rox SYBR kit (Eurogentec). Fluorescence intensity was recorded using a CFX96 Real-Time PCR Detection System (Bio-Rad) and the data analyzed using the ΔΔCt method of analysis. Reference gene 18s was used to normalize the expression level of the gene of interest as previously described (Pfaffl et al., 2001). The selected forward and reverse primer sequences are listed in Table 1. Statistical analyses were performed using GraphPad PRISM 5.0 (La Jolla). Data were analyzed for normal distribution using Shapiro-Wilk test. Non-parametric one-way Anova (n = 4-6) was used to determine transcripts expression level. Primers were designed using Primer 3 software from gene sequences obtained from Genebank. Primer specificity was determined using a BLAST search. For the list of primers used, refer to supplementary table 1.

### Histological staining and analysis

*Tibialis anterior* muscles were collected, embedded in tragacanth gum, and quickly frozen in isopentane cooled in liquid nitrogen. Cross-sections (10μm thick) were obtained from the middle portion of frozen muscles and processed for histological, immunohistochemical, enzymohistochemical analyses according to standard protocols. The fibre cross-sectional area and the number of centrally nucleated fibers were determined using Laminin and Dapi-stained sections. Fluorescence microscopy and transmission microscopy were performed using Axioimager Z1 microscope with CP Achromat 5x/0.12, 10x/0.3 Ph1, or 20x/0.5 Plan NeoFluar objectives (Zeiss). Images were captured using a charge-coupled device monochrome camera (Coolsnap HQ, Photometrics) or color camera (Coolsnap colour) and MetaMorph software. For all imaging, exposure settings were identical between compared samples. Fiber number and size, central nuclei and peripheral myonuclei were calculated using ImageJ software.

### Quantification methods for myonuclei spreading in myotubes

Quantifications in immature myotubes were assessed using an homemade analysis tool. An image analysis performed in ImageJ® software is combined with a statistical analysis in RStudio® software. This provides quantifications of parameters, ranked by myonuclei content per myotubes, regarding phenotype of myotubes (area, length) and their respective myonuclei positioning compare to centroid of myotubes (DMcM).

MSG diagrams were obtained through the normalization of lengths of all analyzed myotubes (independently to their myonuclei content) to 100%. White lines represent myonuclei density curves assessing the statistical frequency for myonuclei positioning along myotubes. Each color group reflects statistical estimation of myonuclei clustering along myotubes.

## Acknowledgements

We thank the Penn Vector Core, Gene Therapy Program (University of Pennsylvania, Philadelphia, US) for providing pAAV1 plasmid (p0001), and Sofia Benkhelifa-Ziyyat for AAV production, the Imaging facilities of Lyon, PLATIM and CICLE, the animal facility of Lyon, PBES, and particularly, Christophe Chamot and Claire Burny for building the macros in ImageJ and R-Studio.

## Author contributions

Conceptualization, A.G., M.B. and V.G.; Methodology, A.G., N.C., M.B., A.G., A-C.D.and V.G. Formal Analysis, A.G., M.B., A.G., A-C.D., D.A., E.C. and V.G. Investigation, G., E.C., M.B., N.C., C.K-R, A.J., A.G., A-C.D, D.A., N-B. R., M-T. B., L.J., M.B. and V.G. Writing – Original Draft, V.G., V.B., and M.B. Funding Acquisition V.G.

---

**Supplementary Table 1.**
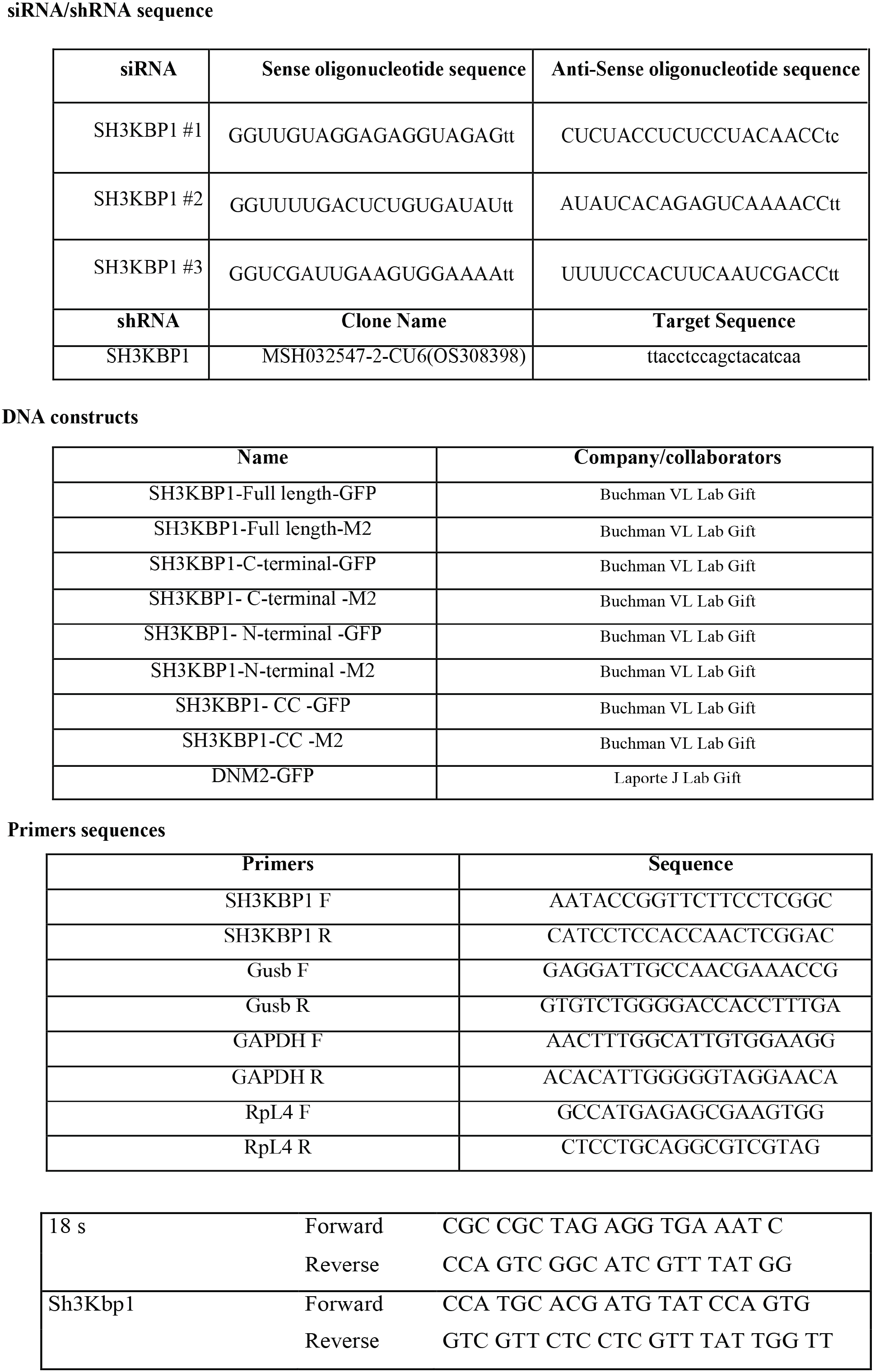

**Supplementary Table 2.**

| Antibody or Dye | Species and Utility | Dilution Factor | Manufacturer and Reference |
| --- | --- | --- | --- |
| Anti-SH3KBP1 | Rabbit | 1:100 | Sigma- HPA003355 |
| Anti-SH3KBP1 (Cin85 H-300) | Rabbit | 1:1000 | Santa Cruz, sc-48746 |
| Anti-SH3KBP1<br>(SETA/Cin85/Ruk/SH3KBP1 Antibody<br>clone 179.1.E1) 05-731 | Mouse | 1:1000 | Merck 05-731 |
| Anti-α-Tubulin | Mouse - primary | 1:1000 | Sigma<br>T6074-200uL |
| Anti-myogenin (F5D) | Mouse | 1:100 | Santa Cruz sc-12732 |
| Anti-RCAS1 (D2B6N) | Rabbit | 1:200 | Cell Signaling #12290 |
| Anti-ERP72 (<u>D70D12</u>) | Rabbit | 1:100 | Cell Signaling #5033 |
| Anti-laminin | Rat | 1:200 | Sigma |
| Anti-GFP (3H9) | Rat | 1:1000 | Chromotek |
| Anti-Flag M2 | Mouse | 1:1000 | Sigma F1804 |
| Anti-Calnexin (C5C9) | Rabbit | 1:100 | Cell Signaling #2679 |
| Anti-LC3II/LC3I | Rabbit - primary | 1:250 | Sigma L7543 |
| Anti-Myosin skeletal slow | Mouse | 1:100 | Sigma M8421 |
| MF20 | Mouse-primary | 1:10 | DSHB |
| Anti-DHPR | Rabbit | 1:10 | DSHB (IIID5E1) |
| Dapi-brilliant blue | Dye | 1:50000 | Thermo Fisher Scientifics<br>D1306 |
| Phalloidin-Alexa Fluor 647 | Dye | 1:100 | Thermo Fisher Scientifics<br>A22287 |
| Anti-mouse-HRP | Goat - secondary | 1:3000 | Invitrogen<br>62-6520 |
| Anti-rabbit-HRP | Goat -<br>secondary | 1:3000 | Invitrogen<br>65-6120 |
| Anti-rat-Alexa Fluor 488 | Donkey -<br>secondary | 1:500 | Thermo Fisher Scientifics<br>A-21208 |
| Anti-rabbit-Alexa Fluor 647 | Goat -<br>secondary | 1:500 | Thermo Fisher Scientifics<br>A-21245 |
| Anti-mouse-Alexa Fluor 647 | Goat -<br>secondary | 1:500 | Thermo Fisher Scientifics<br>A-21240 |

